# Modeling the transmission and vaccination strategy for porcine reproductive and respiratory syndrome virus

**DOI:** 10.1101/2020.05.23.112946

**Authors:** Jason A. Galvis, Joaquin M. Prada, Cesar A. Corzo, Gustavo Machado

## Abstract

Many aspects of the porcine reproductive and respiratory syndrome virus (PRRSV) between-farm transmission dynamics have been investigated, but uncertainty remains about the significance of farm type and different transmission routes on PRRSV spread. We developed a stochastic epidemiological model calibrated on weekly PRRSV outbreaks accounting for the population dynamics in different pig production phases, breeding herds, gilt development units, nurseries, and finisher farms, of three hog producer companies. Our model accounted for indirect contacts by the close distance between farms (local transmission), between-farm animal movements (pig flow), and reinfection of sow farms (re-break). The fitted model was used to examine the effectiveness of vaccination strategies and complementary interventions such as enhanced PRRSV detection and vaccination delays and forecast the spatial distribution of PRRSV outbreak. The results of our analysis indicated that for sow farms, 59% of the simulated infections were related to local transmission (e.g. airborne, feed deliveries, shared equipment) whereas 36% and 5% were related to animal movements and re-break, respectively. For nursery farms, 80% of infections were related to animal movements and 20% to local transmission; while at finisher farms it was split between local transmission and animal movements. Assuming that the current vaccines are 1% effective in mitigating between-farm PRRSV transmission, weaned pigs vaccination would reduce the incidence of PRRSV outbreaks by 2%, indeed under any scenario vaccination alone was insufficient for completely controlling PRRSV spread. Our results also showed that intensifying PRRSV detection and/or vaccination pigs at placement increased the effectiveness of all simulated vaccination strategies. Our model reproduced the incidence and PRRSV spatial distribution; therefore, this model could also be used to map current and future farms at-risk. Finally, this model could be a useful tool for veterinarians, allowing them to identify the effect of transmission routes and different vaccination interventions to control PRRSV spread.

## Introduction

Porcine reproductive and respiratory syndrome virus (PRRSV) is a widespread swine disease in North America (Hermann et al., 2007; Arruda et al., 2018; Sanhueza et al., 2019). PRRSV is seasonal and shown significant epidemic peak during cold months (Perez et al., 2019; Trevisan et al., 2020), causing major economic losses (Holtkamp et al., 2013). PRRSV incidence has decreased in the last couple of years, certainly as a result of biosecurity improvements (Donovan, 2019; Miller, 2019; Thomas, 2019).

In the last two decades, epidemiological studies have described the between-farm PRRSV transmission dynamics (Dee et al., 2003a, 2004; Rosendal et al., 2014; Thakur, Revie et al., 2015; Jara et al., 2020; Andraud and Rose, 2020). Direct contact among farms through pig transportation together with indirect contacts often defined as local spread or local transmission, (e.g. airborne, feed deliveries, shared equipment) have been identified as key routes of swine bacteria and virus dispersal (Velasova et al., 2012; Lee et al., 2017; VanderWaal et al., 2018; Silva et al., 2019; Bastard et al., 2020; Salines et al., 2020). Previous experimental and observational studies have reported frequent PRRSV reinfection events on stable breeding farms (i.e., farrow-to-wean), in this paper we will referred to re-break events (Pileri and Mateu, 2016). Even though there is no clear consensus in the definition of what constitutes a re-break, the rebound of PRRSV could be related to the continued viral shedding within groups of pigs and/or reduced cleaning and disinfection procedures among other unknown factors and dynamics (Bierk et al., 2001; Arruda et al., 2016). Despite the initial findings, the quantification of PRRSV transmission among multiple farm types (e.g., from finisher farms to farrow-to-wean), remains a major gap in PRRSV epidemiology (Jara et al., 2020; Salines et al., 2020).

Reducing the incidence of PRRSV outbreaks is among long term goals of some North American swine producing companies. Currently, the industry relies heavily on vaccination and biosecurity practices as control and prevention strategies (Silva et al., 2019), with the vaccination of weaned pigs and gilts with modified-live vaccines (MLV) being among the most common (Kroll et al., 2018; Garcia-Morante et al., 2020; Rawal et al., 2020). Several studies have evaluated the efficacy of vaccines in increasing the immunity level and reducing shedding magnitude and duration (Pileri et al., 2015; Chase-Topping et al., 2020); however the absence of studies quantifying the effect of vaccination programs on between-farm PRRSV transmission, has limited our capacity to investigate the effectiveness of current vaccination strategies and the development of new, or the enhancement of existing PRRSV vaccination protocols (Andraud and Rose, 2020).

Existing disease spread models have explored the dynamics of between-farm PRRSV transmission (Thakur, Revie et al., 2015; Thakur, Sanchez et al., 2015); a limited number of these studies have investigated the deployment of control strategies on PRRSV outbreak incidence (Bitsouni et al., 2019). To our knowledge no modeling framework has accounted for the spatio-temporal spread dynamics of PRRSV in an epidemiological model. We developed an original mechanistic model that to disentangle the relevance of transmission routes and serve as a framework to test control strategies *in silico* (Brown and Bevins, 2018; Jurado et al., 2019). The current epidemiological model is built to track the between-farm PRRSV spread routes among farm types, breeding herds, gilt development units and growing pigs. The model outputs recorded i) the relative contribution of the between-farm direct contacts by the movement of animals, local transmission by indirect contacts governed mainly by the proximity among farms and exclusively for sow farm we also estimated the probability of re-breaks; ii) the weekly number of new PRRSV outbreaks, iii) the effect of different vaccination strategies in combination with intensified PRRSV detection and delay in vaccination, and iv) the use of the fitted baseline model to forecast and visualize the spatial distribution of at-risk farm locations.

## Material and Methods

We developed a discrete-time stochastic model to simulate between-farm PRRSV transmission. The model included the between-farm pig movements and indirect contacts by local transmission events as a function of the distance between susceptible and infected farms, named hereafter simply as local transmission. The probability of virus to persist within the sow farm named as re-break event was also considered. The survival probability was included to consider PRRSV accumulation in the environment in cases where proper cleaning and drying management cannot be performed (Vilalta et al., 2019) and/or driven by a group of breeding age females or piglets that continue to shed the virus and consequently maintain the transmission chain (Pileri and Mateu, 2016). The methods are divided into sections as follows: i) description of the utilized data, animal movements and the PRRSV outbreak incidence and ii) the mathematical modelling infrastructure including the effectiveness of simulated vaccination strategies on PRRSV transmission and mapping at-risk farms.

### Data sources

Data from a total of 2,294 farms from three U.S. non-related pig production systems (identified as A, B, and C) were obtained from the Morrison Swine Health Monitoring Project (MSHMP) project (Perez et al., 2019). Data from each farm included their national premises identification number, production type (sow [which included farrow, farrow-to-wean and farrow-to-feeder farms]), nursery, finisher [which included wean-to-feeder, wean-to-finish, feeder-to-finish], gilt development unit (GDU), isolation and boar stud, pig spaces per farm, and geographic coordinates.

The between-farm pig flow were used to reconstruct directed weekly contact networks between June 1^st^, 2019 and December 5^th^, 2019. Each pig movement included date, farm of origin and destination, number of pigs transported, and purpose of movement (e.g., weaning). Multiple movement between two farms in the same week were considered as one unique contact and the number of animals moved was summed. Missing movement data (e.g., either farm location, production type, number of animals moved, the farm of origin, or destination) were excluded prior to the analysis. The model included movements between 92% of the farms in the studied region, and comprise a total of 19,984 movements.

The weekly number of PRRSV infected farms between January 5^th^, 2009 and December 5^th^, 2019, was compiled from two sources: i) for the outbreaks in sow farms we used the farm list that each company uses to track farms undergoing outbreaks, re-break or elimination, and ii) for the downstream farms (e.g. growing pig farms) we utilized ORF-5 PRRSV sequence from MSHMP to identify outbreaks, this record is used by the veterinarians to monitor PRRSV strain circulation (Perez et al., 2019; Jara et al., 2020). Briefly, a new sequence is often generated when clinical signs are identified or there is a reason to request a sequence (Jara et al, 2020 for more details about sampling and PRRSV dissemination). For sow farms, we classified each PRRSV occurrence as “new outbreak” or “persistence outbreak” based on the reported outbreak dates. PRRSV detection that occurred within the past 41 weeks from a previously reported PRRSV outbreak in the same sow farm was considered a persistence outbreak, otherwise, it was classified as a new outbreak. We based this classification on the average time to sow farms to reach PRRSV stable status (Sanhueza et al., 2019). Similarly, for nurseries, GDUs, and finishers farms, we used general pig production phase duration to count new PRRSV outbreaks, thus a PRRSV sequence in nurseries, GDUs, and finishers farms within 7, 22, and 25 weeks from a previous record, respectively, was considered a persistent outbreak; otherwise it was counted as a new outbreak. These time windows were based on the general production schedule for nurseries, GDUs, and finisher farms under all-in and all-out system. The infected farms were divided in two groups, for the initial setting we utilized historical cases of PRRSV from January 5^th^, 2009 - June 1^st^, 2019, and for model simulation pig movement and PRRSV outbreaks from June 1st - December 5th, 2019.

### Between-farm infection dynamics

In brief, it was a stochastic model in discrete-time with a weekly time-step. We chose a one-week time-step to match the frequency at which infected farms are reported to the MSHMP project (Perez et al., 2019). Three farm heath statuses susceptible to infection (S), undetected infection i.e., infected but not yet tested or absence of clinical signs (U), and detected infection (D) were considered (Figure 1). In this study, we assumed that all sow farms have been vaccinated or exposed to PRRSV previously, thus all sites have the same probability to report a PRRSV outbreak. The latent period is not explicitly modeled, since at animal level we expect that nasal shedding would be present at day 7 post-inoculation (Pileri and Mateu, 2016; Chase-Topping et al., 2020), thus latent is embedded within the weekly time-step. To model the local transmission, we modified a gravity model to account for the effect of vegetation coverage surrounding each farm, previous showed to modulate between-farm transmission dynamics (Jara et al., 2020), see equations 2-4. Because PRRSV can persist undetected for long periods of time, mainly in contaminated environments and/or fomites (Pileri and Mateu, 2016), for sow farms we included a re-break probability, in which any sow farm that had a PRRSV outbreak in the past two year could have a re-break (Perez et al., 2019; Jara et al., 2020). Transmission rates of the between-farm pig movements, local transmission, and the re-break probability are represented hereafter by the beta values *β*net, *β*local, and *β*rebreak, respectively, fitted to the observed number of PRRSV infected farms (details in the model fitting section).

**Figure 1.**
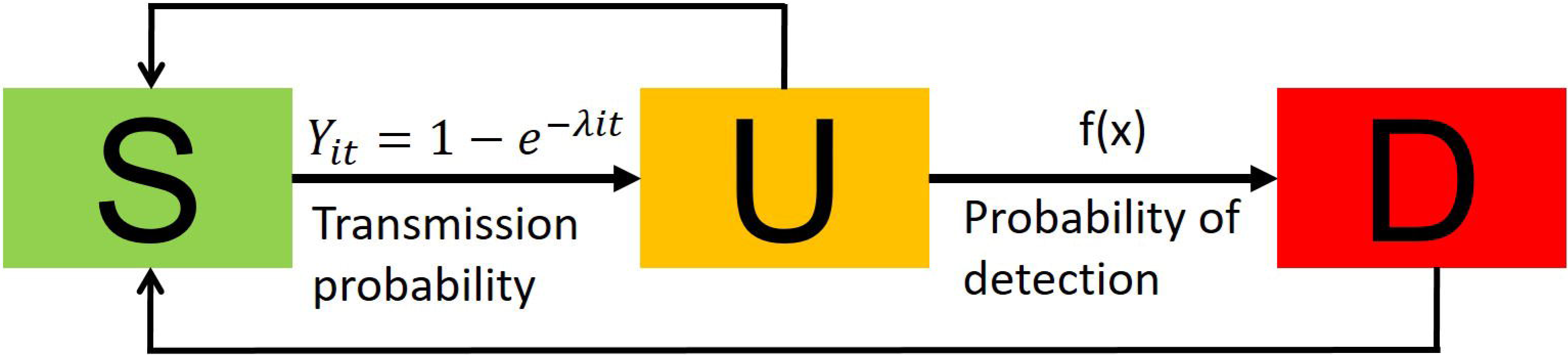
The flow of infection stages in the farm’s modeling framework. Each farm was classified to either state: susceptible (S), infected undetected (U) or infected detected (D).

**Figure 2.**
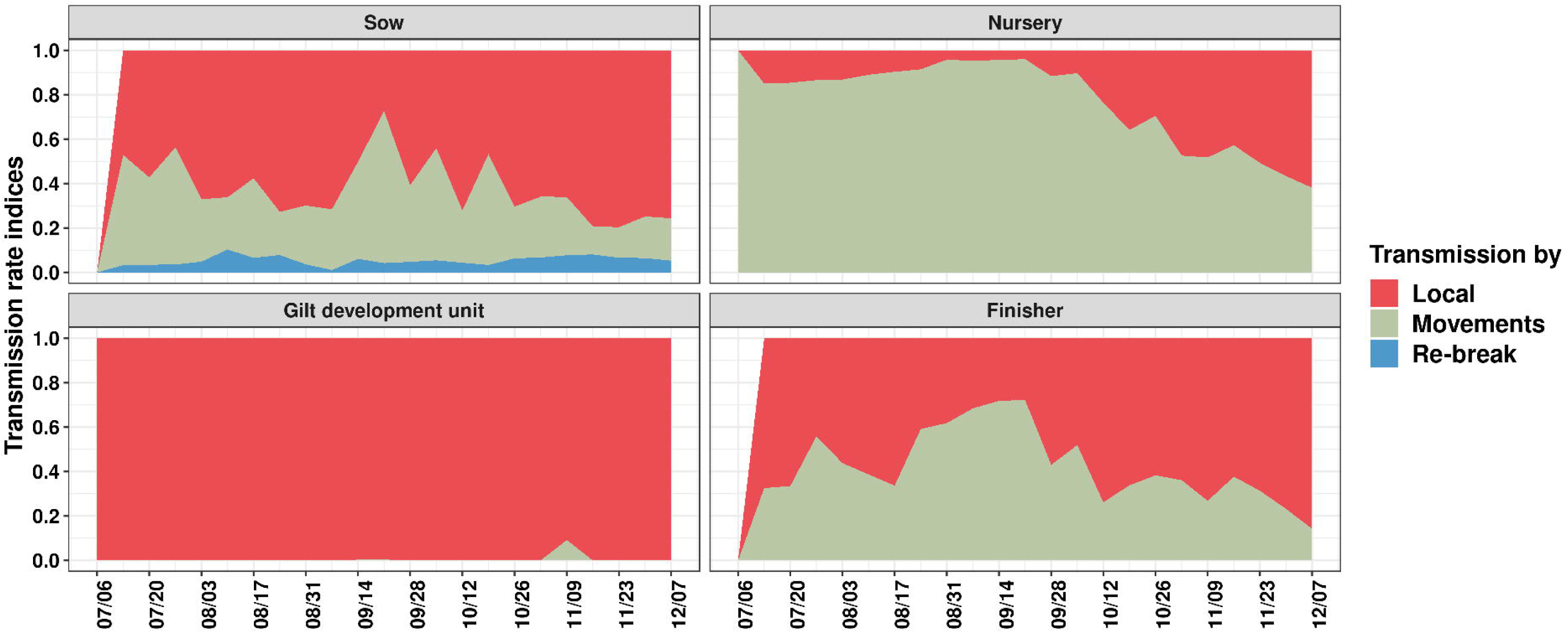
Force of infection for each transmission route of each production type. The y-axis represents the proportion of transmission by transmission routes for each of the simulated weeks (x-axis). Weekly rate indices were calculated by dividing the number of simulated infected farms for each transmission route by the number of simulated infected farms by the three routes combined.

#### Between-farm pig movements

The weekly pig movement network was utilized to identify contacts between susceptible farms *i* and infected undetected farms and detected farms *j*, used to estimate the force of infection driven by the weekly between-farm pig network (**λ**net):

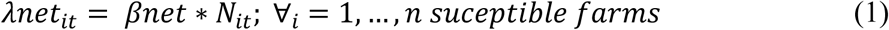

*N*_*i*_ is the number of all infected farms (undetected and detected) sending pigs to susceptible farms *i* at the time step *t* (one week).

#### Local transmission

The local transmission was modeled through a gravity model as follow:

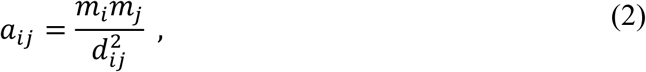

where the attraction force between farms *i* and *j* was directly proportional to the farms’ pig capacity of *m*_*i*_ and *m*_*j*_ divided by the squared Euclidean distance d_ij_^2^ between two farms. Based on a descriptive analysis of the observed outbreaks, the maximum range allowed for the local transmission was set at 35km (*a*_*ij*_ = 0 when *d*_*ij*_> 35 km), *d*_*ij*_ was selected based model performance, it is worth noticing that short or long distances pathogen spread remains a challenge in disease epidemiology in general (Anderson et al., 2017; Machado et al., 2019). In addition, the local transmission was modulated by the enhanced vegetation index (EVI), which was used to quantify vegetation greenness at each farm location, here representing a barrier that reduced local transmission of PRRSV as described elsewhere (Jara et al., 2020). The attraction force, *a*_*ijt*_, between *m*_*i*_ and *m*_*j*_ was modeled as:

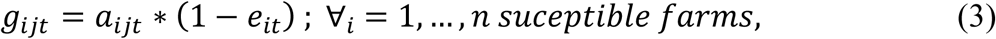

where *g*_*ijt*_ is the attraction force including the barrier effect, in which *e*_*i*_ is the barrier index from the recipient farm *i*, represented on a [0,1] scale (Supplementary section 2 for additional information). Thus, the local infectious force (**λ**local) was defined as:

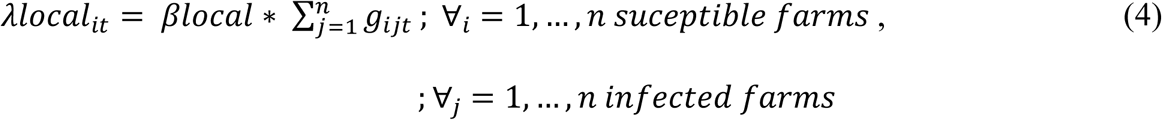

#### The re-break in sow farms

Exclusively for sow farms, a re-break probability was coupled with the other two forces of infection and included as

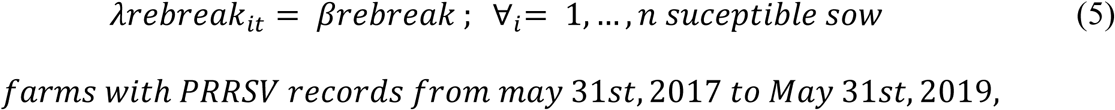

where sow farms that tested positive from May 2017 until May 2019 had equal probability to re-break throughout the simulated period.

### The overall transmission dynamics

The total force of infection (**λ**) was calculated as the sum of all force of infection accounting for the yearly PRRSV seasonality (Jara et al., 2020; Sanhueza et al., 2020),

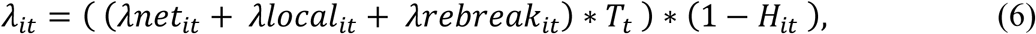

where *λ*_*it*_ is the total force of infection of a susceptible farm *i* and *T*_*t*_ is the seasonality index of time step *t*. The seasonality index values were estimated from the moving averages of the monthly frequency of PRRSV outbreaks between 2015 and 2019, which were then scaled between 0.001 and 1 (Supplementary section 1). The seasonality captured the variability in PRRSV transmission events due to assumed changes in pigs’ immunity and other estimated weather and environmental factors (Dee et al., 2003a, 2003b). Because on-farm biosecurity in sow and GDU farms is expected to play an important role the introduction and spread of new pathogens, we included a biosecurity term (*H*), which modulated *λ*_*it*_. Thus, *H*_*it*_ is defined between 0 and 1, and represents a reduction in the total force of infection. Since no farm information regarding biosecurity measures was available at the moment, we hypothesized that sow and GDU farms without PRRSV historic records were more likely to have enhanced biosecurity measures, thus *H*_*it*_ was applied only for sow and GDU farms without PRRSV outbreaks since 2009 (Table 1), while nurseries and finishers it was assumed to be zero. The *H* value for sow and GDU farms was fitted to the observed number of PRRSV infected farms (details at model fitting section) (Table 1). Finally, we calculated the transmission probability (*Y*_*it*_):

**Table 1.** Transmission parameters used in simulations

| Model parameters | Parameter | Value | Details & references |
| --- | --- | --- | --- |
| Between-farm pig movements (transmission rate) | $\beta_{net}$ | 0.26 | ABC fitting |
| Local transmission (transmission rate) | $\beta_{local}$ | 0.003 | ABC fitting |
| Re-break for sow farms (transmission rate) | $\beta_{rebreak}$ | 0.0019 | ABC fitting |
| On-farm biosecurity levels | $H$ (sow) | 0.47 | ABC fitting |
| | $H$ (GDU) | 0.38 | ABC fitting |
| Detection effectiveness | $L$ (nursery, GDU and finisher) | 0.07 | ABC fitting |
| | $L$ (sow) | 0.95 | Expert opinion |
| PRRSV seasonality | $T$ | Weekly values calculated | (supplementary material section 1) |
| Average time for PRRSV detection | $X_0$ | 4 weeks | Expert opinion |
| Average infectious time for sow farms | - | 41 weeks | (Holtkamp et al., 2011) |

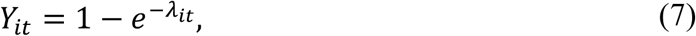

where *Y*_*it*_ is the transition probability from susceptible farm *i* to infected (and undetected) status at time *t*. The detection rate for infected farms (transition from undetected to detected infection status) was assumed to be dependent on the elapsed time of the farms in the infected undetected status and is calculated by a detection probability derived from a logistic function

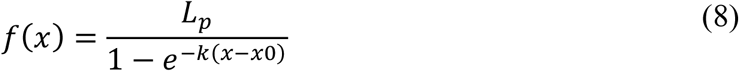

to increase the detection rate at each time step following a sigmoid curve, where *L* is the maximum detection probability for PRRSV outbreaks and its values were defined according to the farm production types *p*. Thus, for sow and GDU farms which undergo periodic testing, we assumed *L* to be high (0.95), while at nurseries and finishers farms (downstream) PRRSV detection is expected to be underestimated, mainly because sampling efforts are much lower than in sow farms. Therefore, in downstream farms *L* was fitted on the PRRSV outbreaks and to the local empirical PRRSV prevalence (details at model fitting section). Here, *X0* is the sigmoid’s midpoint to detect an infected undetected farm (i.e. which was assumed to be the average time for PRRSV detection), defined as four weeks (Neira et al., 2017); *k* is the logistic growth rate, which we assumed to be 0.5, based on the local veterinarian observations in the study region; *X* is the time post-infection (in weeks).

The proportion of recovered sow farms is drawn from a Poisson distribution with a mean of 41 weeks (average time to stability) (Sanhueza et al., 2019). Nursery, GDU, and finisher farms’ transition to susceptible status were driven by pig production scheduling, all-in all-out or by ingoing or outgoing pig movement, whatever come first in the data. Briefly, nurseries, GDU and finisher farms were assumed to become susceptible within 7, 22 and 25 weeks of pig placement, respectively, or when a movement is recorded before reaching the scheduled production phase timeline.

As an initial model setting, farms were considered to be infected if they presented a previous outbreak within a determined number of weeks before June 1^st^, 2019 (initial start point in the simulations) according to their farm type as follows: sow 41, nursery 7, GDU 22, and finisher 25 weeks. All of the remaining farms were assumed to be susceptible.

### Model fitting

We used an Approximate Bayesian Computation (ABC) method based on a modified Sequential Monte Carlo (SMC) rejection algorithm, which allows us to recover the approximate posterior distribution of six model parameters *βnet, βlocal, βrebreak, H* (sow and GDU fitted individually), and *L* (nursery, GDU and finisher farms were fitted to one parameter). A set of draws of these six parameters is denominated a “particle’’ (Beaumont, 2010), and their values were fitted to the observed infected farms for each production type from June 1^st^, 2019, until December 5^th^, 2019.

Although outbreaks were reported at nursery and finisher farms, underdetection is a well-known limitation of PRRSV surveillance in those downstream farms (Velasova et al., 2012). Thus, we used an expected weekly number of nursery and finisher infected farms, derived from an annual prevalence of 30% (personal communications from veterinarians from the production companies we collected the data), and the outbreaks distributed throughout the year according to an estimated seasonality term *T* (Supplementary section 3).

The modified SMC algorithm was a two-step rejection procedure. In the first step, the model was fitted to the weekly number of infected farms by production type using eleven summary statistics (Supplementary Table S1). In the second step, the sensitivity and specificity of the model to predict true risk areas was evaluated through the comparison between simulated and observed infected farm locations falling within square spatial areas of 10 km x 10 km, named hereafter as cells (Supplementary section 3), these final fitting step considered PRRSV outbreaks in sow farms (i.e. true outbreaks reported in our data). The ABC rejection procedure was run to generate 100 accepted particles (N), within the tolerance interval (ϵ) (Supplementary Table S1). We chose the particle with the highest sensitivity in the second step of the fitting procedure for all the subsequent simulations of the different control strategies. Table 1 summarizes the model parameters used.

### The effect of vaccination strategies on PRRSV between-farm transmission

We simulated five vaccinations programs and investigated their impact at reducing the between-farm transmission (Corzo et al., 2010; Azman and Lessler, 2015) (Table 2). We considered a range of vaccine efficacies, here we assumed that mass vaccination reduced the force of infection (the rate at which susceptible to infection farms acquire new infection, Figure 1) by 1%, 2%, 3%, 4%, 5%, 10%, 15%, 20%, 30%, 50%, 80% and 90%. We assumed that vaccine efficacy would have a maximum effect in reducing force of PRRSV between-farm transmission four weeks after vaccination, here it was modeled by a logistic growth curve described before (Kick et al., 2019) (for more details about how *ve* was modeled refer to Supplementary material section 4). Because information about the length of post-vaccination herd-level immunity was not available, we assumed vaccinated pigs maintained protection for 36 weeks, based on expert opinion and previous modelling studies (Evans et al., 2010; Arruda et al., 2016). Therefore, for nurseries, GDUs, and finisher farms, one dose of vaccine protected pigs throughout the whole production phase. The effectiveness of the vaccination strategies by production type was calculated by comparing the reduction in the total number of simulated infected farms for 27 weeks to the counterfactual scenario with no intervention.

**Table 2.**
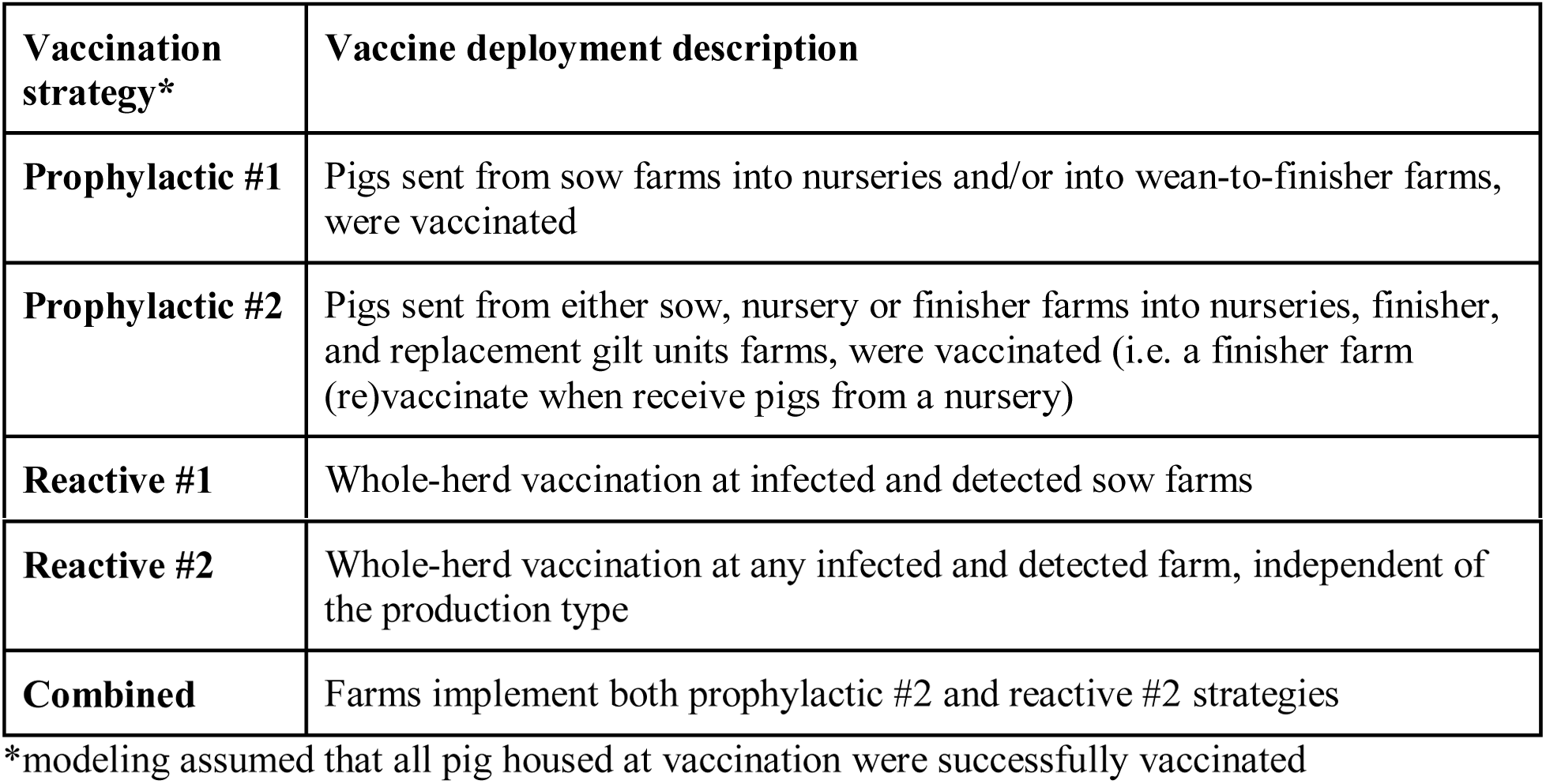
Vaccination strategies

Additionally, we also considered complementary interventions that match more closely current field conditions and practical challenges in terms of vaccination delays and the effect of testing efforts (i.e., decisions about testing finisher farms even if disease is clinical). We examined the impact of i) delaying vaccination for one, two, and three weeks after pig placement, ii) delaying vaccination for one, two, and three weeks post PRRSV detection, iii) enhanced detection scenarios that assumed detection was able to detect 50% or 95% of PRRSV circulation in nurseries, GDUs, and finisher farms, and iv) the average time to detect was set to be between one and three weeks of the virus introduction. All control scenarios were run with 50 repeats to stabilize distributions of the simulated incidence of infected farms. A multivariate linear regression analysis was used to estimate the contribution of each vaccination strategy, individually for each farm type. The outcome of interest was the proportion of prevented PRRSV outbreaks as a function of all complementary interventions while accounting for the range of vaccine efficacies (Supplementary Tables S2 to S6).

### Model outputs

The model outputs included i) the force of infection for each farm type and transmission route, ii) the weekly number of infected undetected and detected farms, iii) the number of prevented PRRSV infected farms for each vaccination strategy, and iv) maps of the location where PRRSV outbreaks are more likely to occur and farms that have recently been infected (recent introductions). We carried out 1000 runs to estimate the relative contribution of each transmission route. Here we evaluated the number of infected farms resulting from each transmission route individually, which were then divided by the number of simulated infected farms from the three combined routes.

To evaluate the model prediction performance of PRRSV infected farm locations, of all sow farms (319 farms) were grouped into 154 cells (10 km x10 km), in which the number of sow farms by cell had a median of 1 and a maximum of 12 farms (Figure 4). The at-risk PRRSV maps were based on the average number of times farms in a given cell were estimated to be infected and posteriorly detected by the model; these cell values were defined as a risk of PRRSV introduction and detection, and were calculated through 100 simulations. Thus, the model spatial forecasting produced two main outputs: a) the risk of PRRSV introduction, here representing areas with farms recently infected; and b) the risk of PRRSV detection, here representing areas with infected but undetected farms that are more likely to be detected. Therefore, we proposed a weekly mapping forecast for both outputs calibrated on the transmission dynamics and observed infected farms from prior weeks; details about predictive performance are described in supplementary material under sensitivity analysis, and an illustrative example is shown in Figure 5.

**Figure 3.**
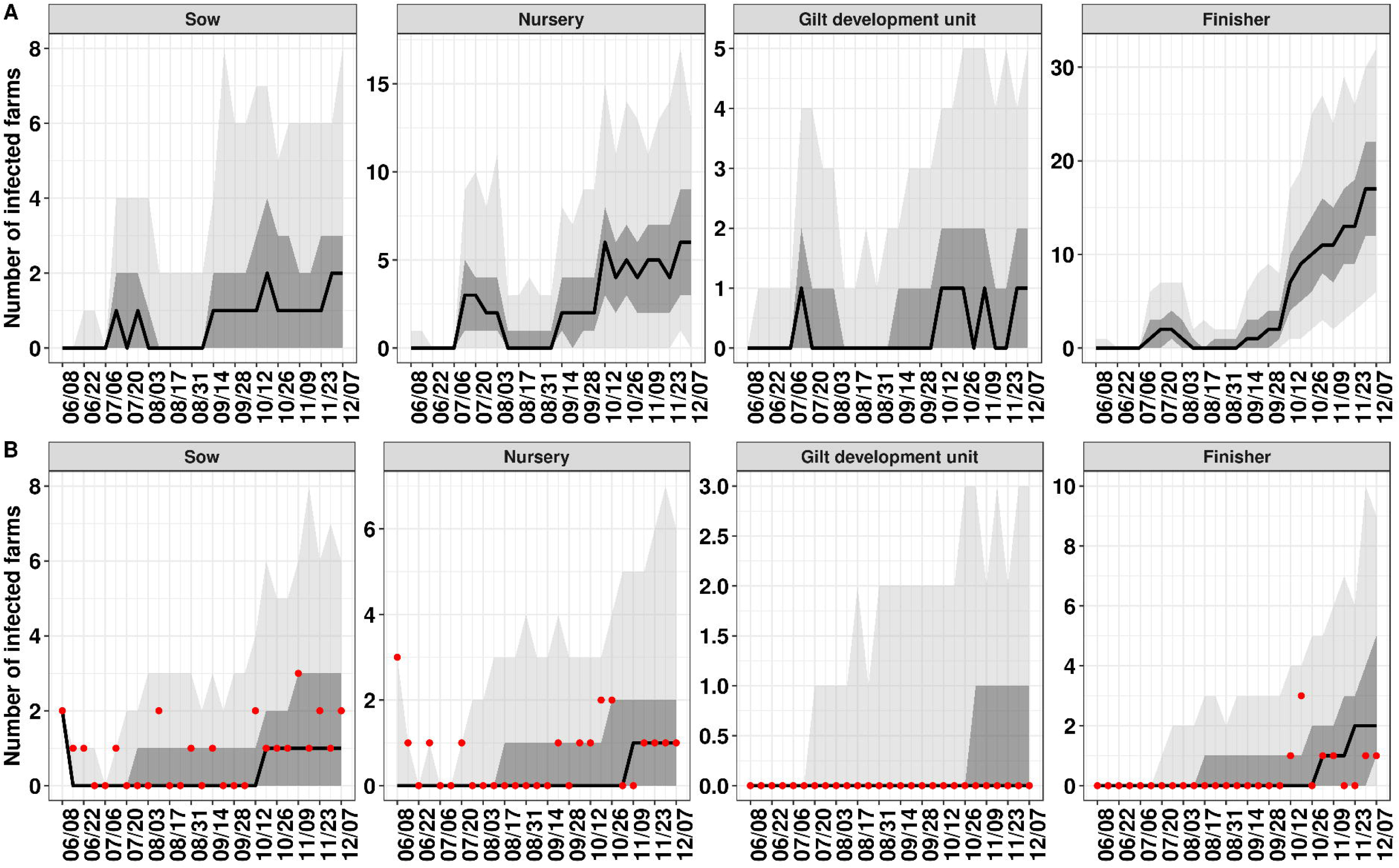
The simulated weekly number of A) infected farms and B) infected and detected farms. The black line represents the median, the dark shade areas represent a 75% credible interval and the light shade areas maximum and minimum generated by the model, and the red dots the frequency of true outbreaks reported in our data. Uncertainty in the estimated model parameters is reflected by 1000 repeated simulations.

**Figure 4.**
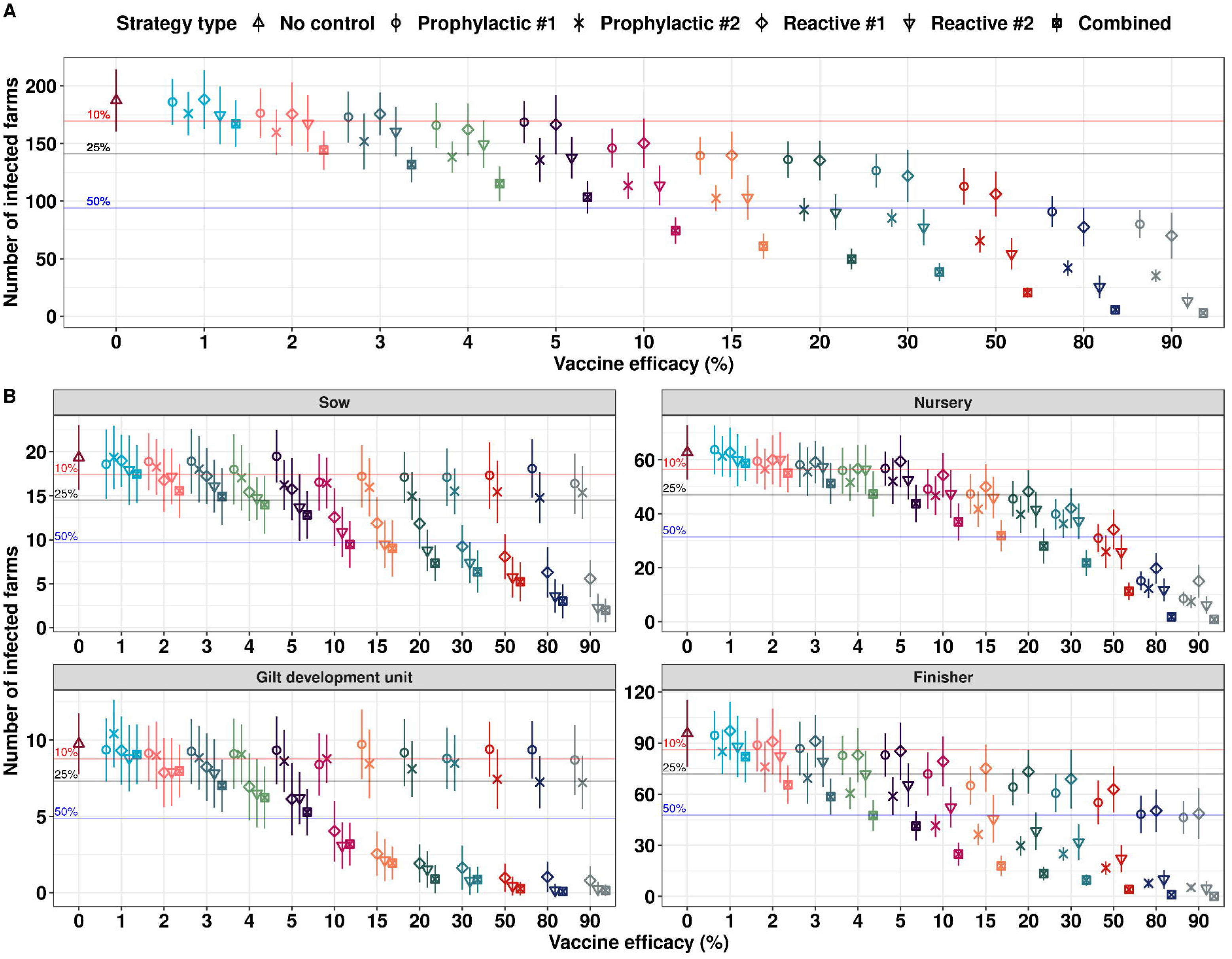
Estimated reduction of PRRSV infected farm trajectories under five vaccine efficacy scenarios. A) The total number of prevented infected farms and B) the number of prevented infected farms depicted by production type. The y-axis represents the number of infected farms within the 27 weeks and the x-axis the simulated vaccine efficacies. Among the prophylactics strategies farms were vaccinated in the same week of receiving a pig movement and among the reactive strategies vaccination occurred in the same week a farm reported an outbreak. The line plot represents one upper and lower standard deviation and the median value represented as the symbol in the center of the line, values calculated from 50 simulations by each vaccination strategy.

**Figure 5.**
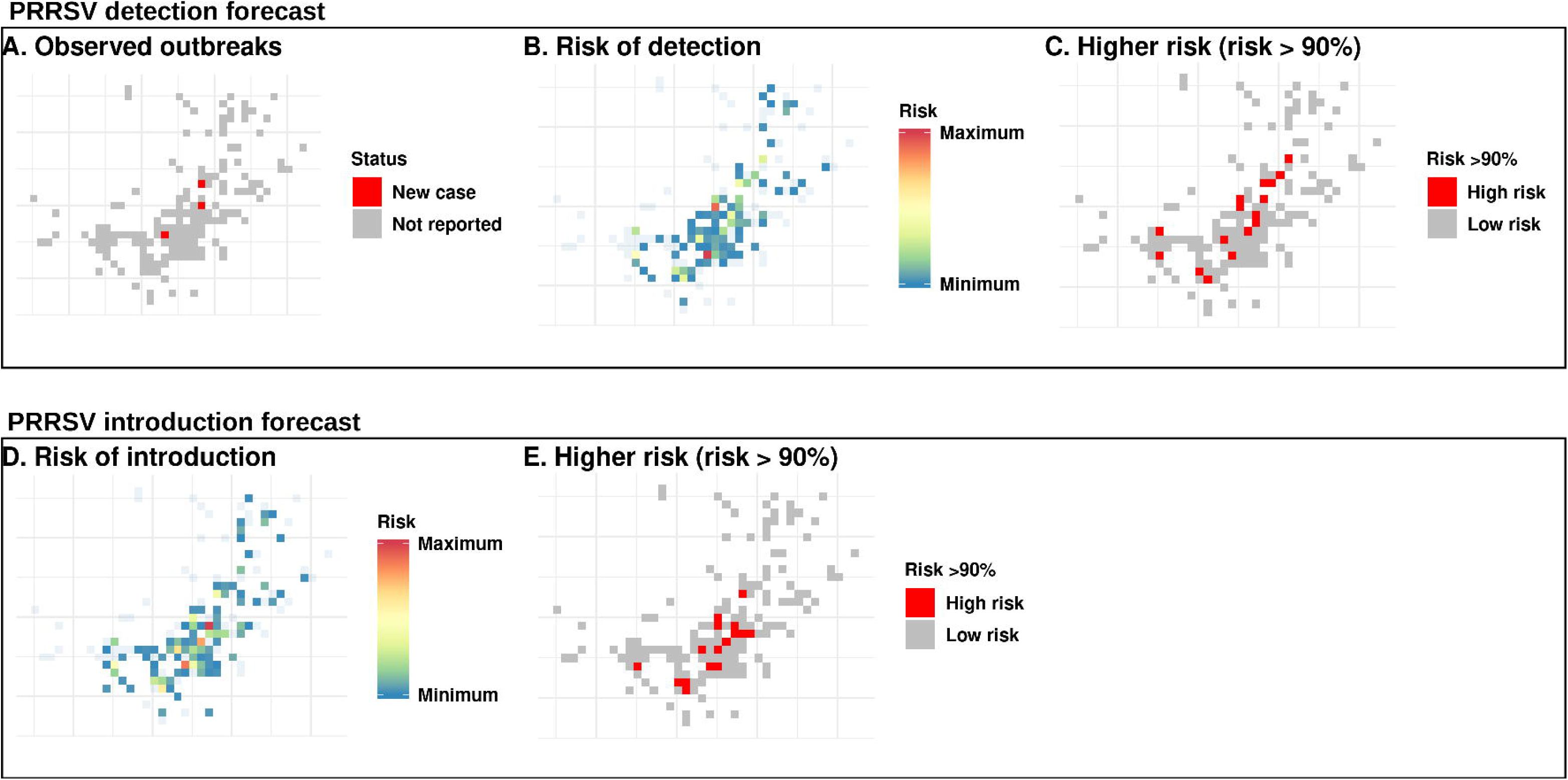
Predicted spatial distribution of PRRSV in sow farms for one target week (November 03-09, 2019). A) cells with observed sow farms infected; B) estimated risk of PRRSV detection over 100 model simulations; C) cells with risk for PRRSV detection above the 90th percentile; D) estimated risk of PRRSV recent introduction; and E) cells with risk for PRRSV introduction above the 90th percentile.

## 3. Results

In Table 3 the number of susceptible and infected farms are summarized. The dataset used to calibrate the model parameters included 48 infected farms reported between June 1^st^ 2019, and December 5^th^ 2019, 27 weeks. Isolation farms and boar studs had no PRRSV detection.

**Table 3.** The summary of susceptible and infected farms by production type from 2009 to 2019.

| **Production type** | **Number of susceptible farms** | **Number of infected farms 2009-2019** | **Number of infected farms Jun-Dec 2019** |
| --- | --- | --- | --- |
| Sow | 319 (14%) | 1070 | 23 |
| Gilt development unit | 187 (8.2%) | 50 | 0 |
| Nursery | 468 (20.4%) | 648 | 17 |
| Finisher | 1272 (55.4%) | 537 | 8 |
| Isolation | 33 (1.4%) | 0 | 0 |
| Boar stud | 15 (0.6%) | 0 | 0 |

### Contribution of each transmission route to PRRSV spread

The results shown here are derived from the fitted particle with higher precision capturing the spatial distribution of infected farms (Table 1). Among the three transmission routes, on average 59% of the simulated infections in sow farms were related to local transmission, and 36% and 5% were related with animal movements and re-break, respectively (Figure 2). For nursery farms, 80% of the simulated infections were related to animal movements and 20% to local transmission, while for finisher, movements and local transmission were similarly important with 47% and 53% of the transmissions, respectively. Finally, GDU, 99% of the infected farms were associated with local transmission.

### The estimated number of new PRRSV infected farms

The estimated average number of new infected farms in the simulated period was 189 (95% CI: 188-191), corresponding to 20 (95% CI: 19-20) newly infected sow farms, 63 (95% CI: 63-64) nursery farms, 10 (95% CI 9-10) GDU farms and 96 (95% CI: 95-97) finisher farms (Figure 3A). There was a good agreement between the weekly observed outbreaks and simulated detected farms (Figure 3B). The model inferred that at the end of the 27 weeks on average 153 of the 189 infected farms would not be detected, in which 81% of the infected sow farms were detected, while a much lower proportion of detection was estimated for GDU 24%, nurseries 11% and for finisher 11% (Figure 3).

### The effectiveness of vaccination strategies

The total number of PRRSV infected farms declined as *vaccine efficacy* increased (Figure 5, and Supplementary Tables S2-S6). A *ve* of 1% was sufficient to reduce on average at least 2% of the total number of infected farms independently of farm types. The combined vaccination strategy at vaccine efficacy of 1% reduced the average number of infected farms by 12%, likewise at *ve*=1% prophylactic #2 and reactive #2 reduced infected farms by 7%, while prophylactic #1 and reactive #1 prevented between 2% and 1%, respectively. Among the five vaccination strategies, the combined strategy showed to be most effective (Figure 4-A), i.e., at *ve = 5%*, the average number of infected farms would be reduced by 46%, which means four times more effective than the prophylactic #1 and reactive #1, and almost two times more effective than prophylactic #2 and reactive #2. The second most efficient strategies were the prophylactic #2 and reactive #2, which showed similar results at *ve* < 20% (Figure 4-A), however reactive #2 performed better with *ve* above 20%. Finally, the prophylactic #1 and reactive #1 strategies marginally reduced PRRSV spread events (Figure 4-A, Supplementary Tables S2-S6).

In addition, in Figure 5-B the simulated vaccination strategies are showed by farm type. Prophylactic #2 showed to work best for finisher farms, while sow, GDU, nursery farms were less likely to be infected under reactive #2 (Figure 4-B). It is worth noting that the combined and reactive #2 strategies rapidly reduced the number of new PRRSV outbreaks in sow and GDU farms, reducing the number of infected farms by 25% with vaccine efficacies of 5% (Figure 4-B).

Supplementary Tables S2-S6 summarizes the results from the multivariate regression analysis, which depicted the effect of delays in the implementation of vaccine interventions and improvements in the PRRSV detections. The effect of complementary interventions ranged from by 0.1% and 4.9%. However, the performance of these complementary interventions depended on the vaccination strategies and the farm type (Supplementary table S2-S6). The most effective interventions was to vaccinate pigs at the same week of placement and increasing the probability of PRRSV detection in downstream farms (Supplementary Tables S2-S6). Overall most complementary intervention showed to have a better impact on PRRSV spread when deployed to finisher and nurseries farms.

### The spatial-temporal risk of PRRSV spread

The calibrated model was used to map the PRRSV spatial risk patterns. The direct comparison of predicted infected cells with the true infected farms weekly, was used to evaluate the model’s predictive ability. Under the an optimal threshold for disease detection, explained in details in supplementary Figure S3, the model predicted correctly infected sow farms with a median sensitivity of 62%. Figure 5 shows the main spatial features of the modelling forecast. Here we defined high-risk cells were all values with risk above the 90^th^ percentile; this percentile value was chosen as a rigorous threshold in order to evaluate the model capabilities to identify locations with high infection risks with the least number of cells (Supplementary Figure S4). In Figure 5-A, we show a map with the true outbreaks reported in our data, Figure 5-B shows the predicted risk of infection, and Figure 5-C shows the cells with risk values above the 90^th^ percentile threshold. In this example two of the three truly infected cells were correctly predicted (Supplementary Figure S4). In addition, the model was also used to estimate the risk of a recent PRRSV introduction (Figure 5-D and 5-E), thus the identified cells more likely to have at least one infected farms which has not yet been identified as positive.

## 4. Discussion

We demonstrated that the dynamics of between-farm PRRSV transmission was strongly modulated by farm types. Our results showed that local transmission generated the majority of the infections in sow (that include farrow, farrow-to-wean and farrow-to-feeder farms) and GDU farms; however, nurseries were more likely infected through animal movements as expected, while for finisher farms (which included wean-to-feeder, wean-to-finish, feeder-to-finish farms) both routes were equally important. Quantifying the routes of transmission can be used in the development of new disease control strategies targeting the dominant transmission routes at each specific farm type (Andraud et al., 2019; Machado et al., 2019). With regards to control strategies, we evaluated the effect of vaccination, enhanced detection and diagnostic performance on the number of prevented PRRSV infected farms. The vaccination of pigs at placement (prophylactic #2) combined with mass-vaccination during PRRSV outbreak (reactive #2), showed to be the most efficient strategy. It is worth noting that the performance of vaccination strategies with the same efficacy varied among farm types. The best vaccination approach for sow and GDU farms was reactive #2. For nursery, the vaccination of weaned pigs (prophylactic #1) was nearly as good as (re)vaccinating finishers and gilts at placement (prophylactic #2). Similarly, prophylactic #2 was highly efficient to prevent infection in finisher farms, and implement vaccination within one week of placement was the best complementary strategy. We also showed that the control of PRRSV spread can be optimized if diagnostic in downstream farms is enhanced and if vaccination is deployed early on PRRSV infection (Henao-Diaz et al., 2020). Finally, based on our baseline model we present two mapping results, a forecasting map showing the most likely infected farms and a map showing at-risk locations in which the risk that PRRSV has been introduced within an average of four weeks was estimated to be high. Ultimately, this can help the identification of farms where interventions would be more effective at reducing the between-farm transmission (Stevenson et al., 1993; Bilodeau et al., 1994; Hayama et al., 2013, 2020).

### Contribution of transmission routes

The important contribution of the local transmission route in spreading PRRSV into sow, GDU and finisher farms was consistent with previous studies, which demonstrated that farms located close to infected sites were more likely to report PRRSV outbreaks (Velasova et al., 2012; Phoo-ngurn et al., 2019; Silva et al., 2019; Jara et al., 2020), while others suggested that contact networks was the preferred transmission route (Amirpour Haredasht et al., 2017; Lee et al., 2017; Bastard et al., 2020; VanderWaal et al., 2020). Our approach to model the local PRRSV spread was unique in that we considered the effect of vegetation around each farm location as a physical barrier against mainly airborne transmission (Van Ryswyk et al., 2019; Jara et al., 2020). Consequently, in our modelling framework, farms at a location with a high vegetation index were less likely to become infected by local transmission route (Jara et al., 2020).

Interestingly, finisher farms transmission rates of finisher farms were less related to the in-going movements when compared with nurseries. This could be attributed to several factors that likely facilitated the between finisher farms local transmission, including high regional finisher farm PRRSV prevalence (personal communication, local veterinarians) and the disproportionate number of susceptible finisher farms. Descriptively, the median number of finisher farms in a 5 km radius around each finisher farms (hereafter defined as a neighborhood) was nine, which was higher than the median of six finishers around nursery farms (Supplementary Figure S5). It is worth considering that the number of pigs flowing into the neighborhoods densely populated with finisher farms is expected to be higher, which could also play an important role in sustaining spread at a local scale due to the regular reintroduction of the virus (Otake et al., 2010; Machado et al., 2019; Ezanno et al., 2020). Neighborhood contacts generally occur in a continuous manner over rather long periods of time (Ezanno et al., 2020), which rise indirect contacts (e.g., airborne) which would vary over time. Similarly, the intense local transmission into sow and GDU farms may be equally associated with a high number of farms in those neighborhoods, especially in neighborhoods heavily populated by farms with known restricted biosecurity, such as finisher farms (Supplementary Figure S5). The results of our local transmission route could be also associated with indirect contacts via vectors and fomites (e.g. farm visitors, other animals, insects, and contaminated transport-tools) (Pitkin, Deen and Dee, 2009; Pitkin, Deen, Otake et al., 2009; Reiner et al., 2009; Lowe et al., 2014). In future work it would be pivotal to include indirect contact networks such as feed deliveries and mortality management through dead pickups (Porphyre et al., 2020), which would allow PRRSV to spread throughout multilayer networks, for example. This will likely shed light on other routes involved in PRRSV spread (Dee et al., 2020).

As described elsewhere, the number of in-going movements into sow and GDU farms have not been associated with the introduction of pathogens (Lee et al., 2017; Bastard et al., 2020), indeed, poorly connected sow and GDU farms are expected to be less infected through receiving infected. In our study, this could be one of the reasons why pig movements were less relevant for sow and GDU farms than the local transmissions; however, a thorough network analysis is still necessary to fully evaluate this findings. Nursery and finisher farms on the other hand are more prone to be infected by animal movements, previously associated with their network position (Passafaro et al., 2020), as observed in other studies (Lee et al., 2017; Bastard et al., 2020), similarly to the result described here.

This study uniquely considered PRRSV re-break events as a source of infection for sow farms, which appears to be significantly lower in comparison with the local transmission and the pig movement routes. On average, the re-breaks were associated with 5% of the outbreaks in sow farms. Given the assumption and the lack of data to calibrate the re-breaks, it is possible that this transmission route has been underestimated. Indeed, it does not mean that re-breaks are not relevant in the epidemiology of PRRSV. Previous investigations described the relevance of sub-populations which could maintain the virus population shedding for an extended period without being identified (Bierk et al., 2001; Evans et al., 2010; Pileri and Mateu, 2016). Our limited understanding of PRRSV infection dynamics in terms of herd immunity added to the complexity of the population dynamics inside multi-barn pig farms, such as in sow farms that include farrowing and gestation barns and the constant movement of animals, equipment and people across barns is a major gap that needs further analysis before we can disentangle the role of re-breaks in the epidemiology of PRRSV.

### The effectiveness of control strategies

The vast majority of commercial pig producers in the US currently utilize commercial modified live-virus (MLV) vaccines (Pileri and Mateu, 2016; USDA, 2016). The limited effectiveness of vaccines in field conditions (Charerntantanakul, 2012; Roca et al., 2012), highlight the need to estimate the level of protection required to successfully reduce PRRSV between-farm transmission. As describe elsewhere (Linhares et al., 2012), suggested that vaccination can reduce PRRSV shedding rates, a more recent experiment with a European type subtype 1 strain, indicated that the current vaccines have a poor efficacy to reduce between animal transmission (Chase-Topping et al., 2020). Therefore, remain under debate the impact MLV vaccines on the concentration of PRRSV in the environment and the reduction of virus shedding. Thus *ve* for between-farm transmission at 1% may be a fair assumption, however proper experimental studies are needed to quantify it. According to our modeling, a vaccine efficiency of 80% would be necessary to completely prevent new transmission events. Our simulations reinforce the need for improving the current vaccine technologies (Bitsouni et al., 2019), meanwhile non-therapeutic interventions such as enhancement of on-farm biosecurity will continue to be essential in reducing PRRSV spread (Silva et al., 2019).

When comparing the effectiveness of prophylactic strategies #1 and #2 for finisher farms, the former strategy was better. The extra vaccine boost likely reduced the force of infection of both routes, animal movement and local transmission, either by increasing the protection of susceptible farms or by decreasing the force of infection from infected and vaccinated farms. On the other hand, both prophylactic strategies provided limited protection for sow and GDU farms (Figure 2 and 5). An added value of prophylactic #2 is that finisher farms may be the main source of virus to other farm types within the neighborhood. In fact, studies showed that the vaccination of finisher sites resulted in a reduction of the likelihood of airborne transmission and or transmission by fomites due to the reduction of nasal shedding in vaccinated pigs (Linhares et al., 2012; Pileri et al., 2015). It is important to remark that vaccination of either prophylactic program showed to work best in nursery and finisher farms if deployed in the same week of placement when compared with a later vaccination, for example 3 weeks after pigs arrive.

The vaccination of PRRSV positive farms was directly dependent on virus detection and reporting (Perez et al., 2019). Therefore, if all farms with detected PRRSV infection were vaccinated as proposed in the reactive #2 strategy, the number of prevented infected farms was approximately the same as the preventive prophylactic #2 scheme, described above. The addition of nursery, GDU and finisher farms in the vaccination of infected farms (reactive #2) was better than reactive #1, even though the model assumed that PRRSV detection rates in downstream farms was at 7% (Table 1). The main limitation of reactive #2 is related to PRRSV diagnostic in finisher and nursery farms. The reactive #2 strategy could potentially be more effective than the prophylactic #2 in terms the volume of vaccine doses. On the other hand, the prophylactic #2 strategy would be independent from diagnostics costs, and thus implementation could be considered more feasible by the swine industry. Before recommendations can be made, future longitudinal studies need to evaluate the economics of each vaccination strategy taking into consideration the cost to stabilize an infected farm (Linhares et al., 2015; Nathues et al., 2017; Thomann et al., 2020). Until diagnostic techniques become more accessible and easier to perform, the effectiveness of strategies involving the identification of PRRSV in downstream pig operations remains questionable.

### The spatial-temporal risk of PRRSV

Unlike many other modelling frameworks, which calibrate models explicitly to fit on the number of outbreaks over time (Brooks-Pollock et al., 2014; Funk et al., 2018), here we proposed the calibration to both the space- and time-distribution of PRRSV outbreaks (Sumner et al., 2017; VanderWaal et al., 2018). This approach allowed a closer approximation to the weekly incidence and the true infected farms (Widgren et al., 2018), thus our model was able to correctly capture at-risk areas with at least one infected farm (Supplementary Figure S3). It is important to note that the accuracy of the forecast is expected to vary as PRRSV incidence changes along the season (Trevisan et al., 2020); therefore, throughout our weekly simulations, the overall sensitivity ranged from 50% to 73%, at the proposer optimal risk thresholds. Improving the spatial forecast sensitivity to reproduce the PRRSV outbreaks spread an important challenge, this is because fitted models to real data may not capture completely the disease transmission fluctuation, similarly to other studies that have also attempted to model the distribution of *E. Coli* or Porcine Epidemic Diarrhea (VanderWaal et al., 2018; Widgren et al., 2018; Machado et al., 2019). Our study was in agreement with a previous study that also highlighted the challenges to predict PRRSV infection in sow farms, when compared with nurseries, wean-to-finish and finisher (Amirpour Haredasht et al., 2017). The challenges of model fitting could be attributed to the lack of information about other contact networks that connect farms indirectly, such the routes taken by feed deliveries and dead picks, which could explain in a greater detail the spatial dispersion of PRRSV. Recent work reinforced the relevance of the indirect contacts among farms, in which sharing haulage vehicles has significant potential for spreading infectious diseases within the pig sector in Great Britain (Porphyre et al., 2020). Therefore, futures including other indirect contact networks into our model framework, would also likely to improve our forecast accuracy.

The forecast sensitivity of 62% was obtained under the optimal risk threshold; however, the number of cells identified to be at greater risk of PRRSV infection, were on average 55 cells. In order to define a feasible threshold for the identification of at-risk areas, there is a need to not only consider model precision, but also the economics of possible targeted deployment of control strategies. In the example shown in this study (Figure 5), we proposed high risk cells to be those with risk above the 90^th^ percentile, thus on average 16 cells were identified to be at risk. As result the model sensitivity was lower than the overall mean value (Supplementary Figure S3). On the other hand, less being more extract to which cells are at risk, made this approach more useful, and informative for the stakeholders’ response and decision making, which could include the formation of a forecast dashboard to expedite risk communication and guide mode precise interventions (e.g., vaccination). A more uneconomic threshold could be more effective in identifying outbreaks, but it would be necessary to evaluate the capacity of the production companies to intensify testing before these thresholds can be proposed. Our model also identified cells that did not present outbreaks (false positives), which could be associated with the presence of infected not yet detected farms. The swine industry could directly benefit from this approach by testing farms in which the model predicted a recent PRRSV introduction or enhance control strategies i.e., vaccination, enhancement of on-farm biosecurity to specific farms (Brooks-Pollock et al., 2014).

### Limitations and further remarks

Our study has a number of simplifications and limitations starting with the study being conducted in one region where three production companies are present which may not entirely represent what occurs throughout the country. Also, the number of cases reported in downstream farms were counted based in the number of ORF-5 PRRSV sequences, which was likely underestimated, mainly because those sites do not undergo systematically testing. In addition, the fact that individual veterinarians’ together with the diagnostician decide which sample to be sequenced may underestimate viral diversity. However, thus far we believe that the use of ORF-5 PRRSV sequences to capture viral circulation in growing pig farms is still the best alternative to the current PRRSV available databases. In addition, we did not consider the influence of individual viral lineages while modeling PRRSV transmission forces, we assumed that all samples had the same virulence, thus the probability of the between-farm transmission was assumed to be the same (Jara et al., 2020). Modelling individual PRRSV strain dissemination was beyond the scope of this study, but this is likely a potentially important future consideration as part of our active research focus area. In the current modelling framework, we were interested in the between-farm transmission, in which we did not include more detailed dynamics of individual pigs within farms, thus futures studies including within-farm dynamics at animal level will be an important component of future studies (Andraud and Rose, 2020). In addition, the model was calibrated on the incidence across all farm types for the temporal model calibration, while we decided to fit the spatial component to infected sow farms only, largely because PRRSV outbreaks records in sow farms were more consistent allowing for a more confident assessment of model sensitivity. Transmission events not explained by between-farm movements or re-breaks at sow farms were classified in the model as local transmitted, thus future studies will include other movement flows such as feed truck deliveries in order to explain in more details this route of transmission (Porphyre et al., 2020). For the assessment of vaccination strategies, we assumed that vaccinated farms acquire life-long immunity, future studies should also consider the heterologous host responses to vaccination (Ezanno et al., 2020).

## Conclusion

This study establishes a modeling framework to explore between-farm PRRSV transmission, which was used to a) quantify the contribution of transmission routes for individual farm types, b) quantify the number of infected farms prevented by vaccination strategies, and c) forecast current and future PRRSV spatial distribution. Our findings reveal that local transmission is a key driver of PRRSV spread into sow and GDU farms, whereas animal transportation into downstream farms causes the majority of the infections in nurseries. Animal transportation and local transmission are equally responsible for introducing infections into finisher farms. Vaccination alone was insufficient for completely controlling PRRSV spread. Weaned pigs vaccination showed to be effective, however when we assumed that vaccine efficacies were at ≥3% and ≥10%, the model estimated 10% and 25% reduction in PRRSV outbreaks, respectively. Our results also reiterated that the implementation of additional strategies that can reduce PRRSV collateral spread, such as on-farm biosecurity, will continue to play a key role in reducing collateral transmission, especially if those strategies are also applied in finisher farms, from which PRRSV was easily spread via local transmission route. The detailed outputs of our spatially explicit model closely reproduce the empirical spatial distribution of PRRSV infection in sow farms. For this reason, the risk forecasted by the fitted model has the potential to guide targeted control interventions. Ultimately, this study offers producers and swine veterinarian’s greater insight into mechanisms of PRRSV spread between different farm types, and how vaccination programs could more effectively in reducing disease transmission.

## Acknowledgements

The primary funding support of this project is from the Swine Health Information Center under project #19-211. This work was also supported by Critical Agricultural Research and Extension 2019-68008-29910 from the USDA National Institute of Food and Agriculture. This project also fostered international collaboration between Dr. Machado and Dr. Prada laboratory from the travel support provided by the University Global Partnership Network (UGPN) Research Collaboration Fund. The Morrison Swine Health Monitoring Project is a Swine Health Information Center funded project. Authors would like to acknowledge participating systems and veterinarians.

## Authors’ contributions

JAG and GM conceived the study. JAG, JP, and GM participated in the design of the study. CC coordinated the PRRSV data collection via MSHMP. JAG conducted data processing, cleaning, designed the model, simulation scenarios with the assistance of JP and GM. JAG designed the computational analysis. JAG, JP, and GM wrote and edited the manuscript. All authors discussed the results and critically reviewed the manuscript. GM recurred the USDA-NIFA funding.

## Conflict of interest

All authors confirm that there are no conflicts of interest to declare

## Ethical statement

The authors confirm the ethical policies of the journal, as noted on the journal’s author guidelines page. Since this work did not involve animal sampling nor questionnaire data collection by the researchers there was no need for ethics permits.

## Data Availability Statement

The data that support the findings of this study are not publicly available and are protected by confidential agreements, therefore, are not available.

## Notes

### Competing Interest Statement

The authors have declared no competing interest.

