## supplementary for "Modeling the transmission and vaccination strategy for porcine reproductive and respiratory syndrome virus"

**Running title:** The transmission dynamics of PRRS virus

Jason A. Galvis[1], Joaquin M. Prada[2], Cesar Corzo[3], Gustavo Machado[1*]

[1] Department of Population Health and Pathobiology, College of Veterinary Medicine, Raleigh, North Carolina.

[2] School of Veterinary Medicine, Faculty of Health and Medical Sciences, University of Surrey, Guildford, UK

[3] Veterinary Population Medicine Department, College of Veterinary Medicine, University of Minnesota, St Paul, MN, USA

**Section 1: PRRSV seasonality**

By evaluating the monthly number of PRRSV outbreaks from 2009 to 2019, we confirmed the presence of a yearly PRRSV season, further used in the modeling steps(Figure S1).

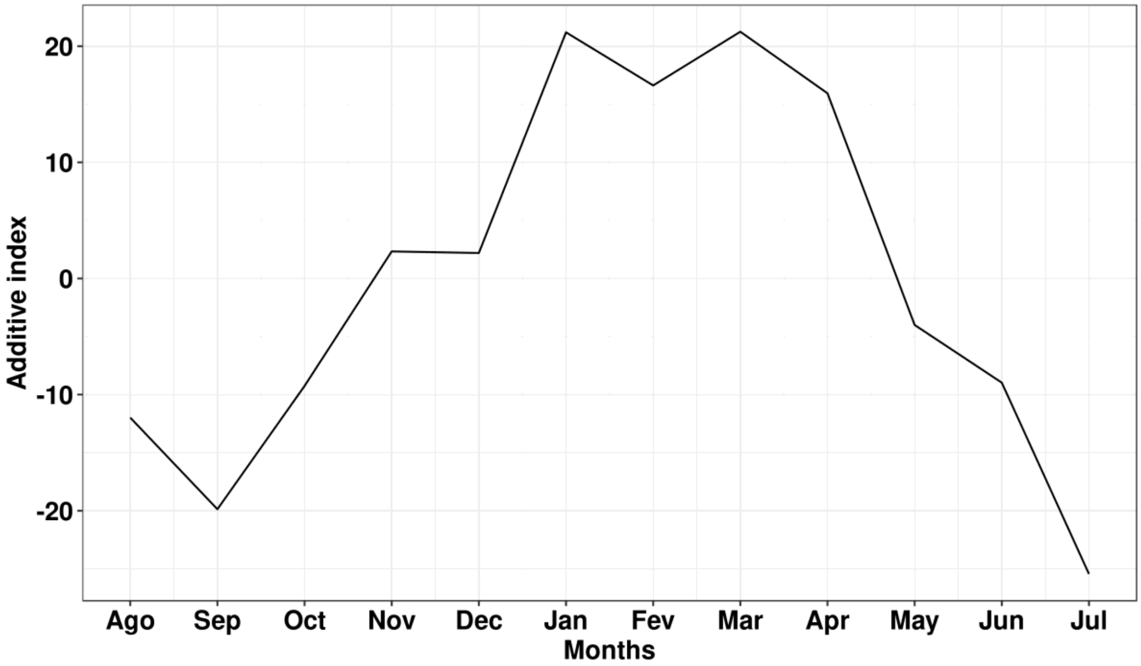


**Figure S1.** Monthly seasonality index calculated from the frequency of PRRSV, calculated by an additive moving average decomposition.

**Section 2: Gravity model with barrier**

To calculate the barrier index (vegetation level, utilized to modulate the probability of local transmission), we used a linear regression, to express PRRSV infected farms between 2018 and 2019 as a function of the Enhanced Vegetation Index (EVI) and yearly seasonality (spring, summer, fall and winter). We found that PRRSV frequency decreased as EVI increased, with a stronger association in winter and spring seasons (Figure S2). Here we used the regression coefficients to predict weekly PRRSV incidence, which then were transformed into parameter *a*, which was scaled into values between [0, 1], later utilized to modulate the local transmission.

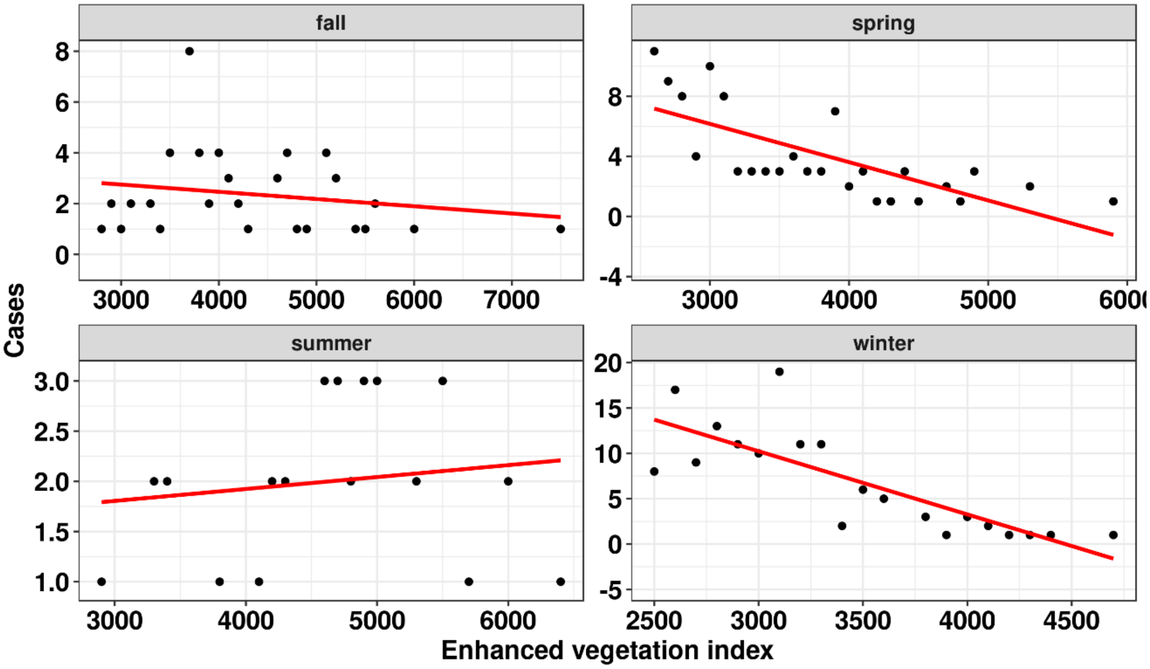


**Figure S2.** Linear regression of PRRSV infected farms. Y-axis the number of PRRSV outbreaks and in the x-axis EVI.

**Section 3: Model fitting, sensitivity, and specificity**

Table S1 describes the summary statistics used in step 1 of the ABC rejection algorithm, where the tolerance interval represents the square error allowed in the simulations. To calculate the frequency of infected farms in nursery and finisher farms considering the yearly PRRSV seasonality (Jara et al., 2020), the empirical prevalence (30%) was distributed to each week according to the seasonal index *T* calculated previously, and the monthly average number of weeks. For example, the expected prevalence for each week in December was calculated as $(0.30/(T_{December}/\Sigma^{n=12}_{m=1}T_m))$/(monthly average number of weeks = 4.34); $\forall_m$=1 (January),...,12 (*December)*.

**Table S1.** Summary statistics used by the Approximate Bayesian Computation Sequential Monte Carlo rejection algorithm.

| **Summary statistic** | **Observed values** | **Tolerance interval** ($\epsilon$) |
|---|---|---|
| Total number of sow farms with detected cases | 23 | 10 |
| The weekly average number of sow farms with detected cases | 0.85 | 0.5 |
| The weekly maximum number of sow farms with detected cases | 3 | 5 |
| Total number of nursery farms with detected cases | 17 | 10 |
| The weekly average number of nursery farms with detected cases | 0.62 | 0.5 |
| The weekly maximum number of nursery farms with detected cases | 3 | 5 |
| Total number of finisher farms with detected cases | 8 | 10 |
| The weekly average number of finisher farms with detected cases | 0.3 | 0.5 |
| The weekly maximum number of finisher farms with detected cases | 3 | 5 |

| | | |
|---|---|---|
| Total number of expected nursery and finisher farms infected | 155 | 100 |
| The weekly average number of expected nursery and finisher farms infected | 5.7 | 5 |
| Maximum number of expected outbreaks in nursery and finisher farms | 20 | 10 |

To assess the model performance, we evaluated the probability to predict cells (10 x 10 km squares) with true infected cells (cells where at least one sow farm outbreak was recorded) at time *t*. Each farm was allocated to a cell in the spatial grid (total of 154 cells in the study area). The risk of each cell was calculate by the average number of times farms within a cell were identified to be infected detected, after 100 simulations. Based on the distribution of the estimated risk values, we utilized a percentiles thresholds (r) approach to determine cells at high and low risk. Where high risk cells were compared with the true infected cells at time *t*. Subsequently we estimate the model sensitivity and specificity, for all thresholds showed in Figure S3, as follows:

$$S_r = TP_r/(TP_r + FN_r)$$

$$E_r = TN_r/(TN_{rt} + FP_r)$$

where true positives (TP) was the subset of cells with observed outbreaks and where the estimated risk was *above* the (r) threshold; false negatives (FN) was the subset of cells with observed outbreaks and where the estimated risk was *below* the (r) threshold; true negative (TN) was the subset of cells without observed outbreaks and the estimated risk was below the *r* threshold; and false positives (FP) was the subset of cells without observed outbreaks and the estimated risk above the *r* threshold. It is worth noting that cells with zero risk were not considered in the sensitivity analysis.

Regarding the model calibration, the sensitivity and specificity, it was calculated for each particle accepted in the step 1 of model fitting (ABC). The particles with sensitivity values ≥45% with r = 85$^{th}$ and ≥50% with r = 75$^{th}$ were accepted in the second step of the model fitting.

The overall sensitivity and specificity of the model for each (r), was based in the average *S* and *E* from (r) values of 1 to 100 by running 1,000 replications. Here, the model showed a maximum medium sensitivity of 67% (minimum 55% and maximum 83%) with $r$ = 40$^{th}$, after that the threshold the model could not correctly predict any additional infected cell and the sensitivity was constant regardless the decreasing of the (r) threshold (Figure S3).

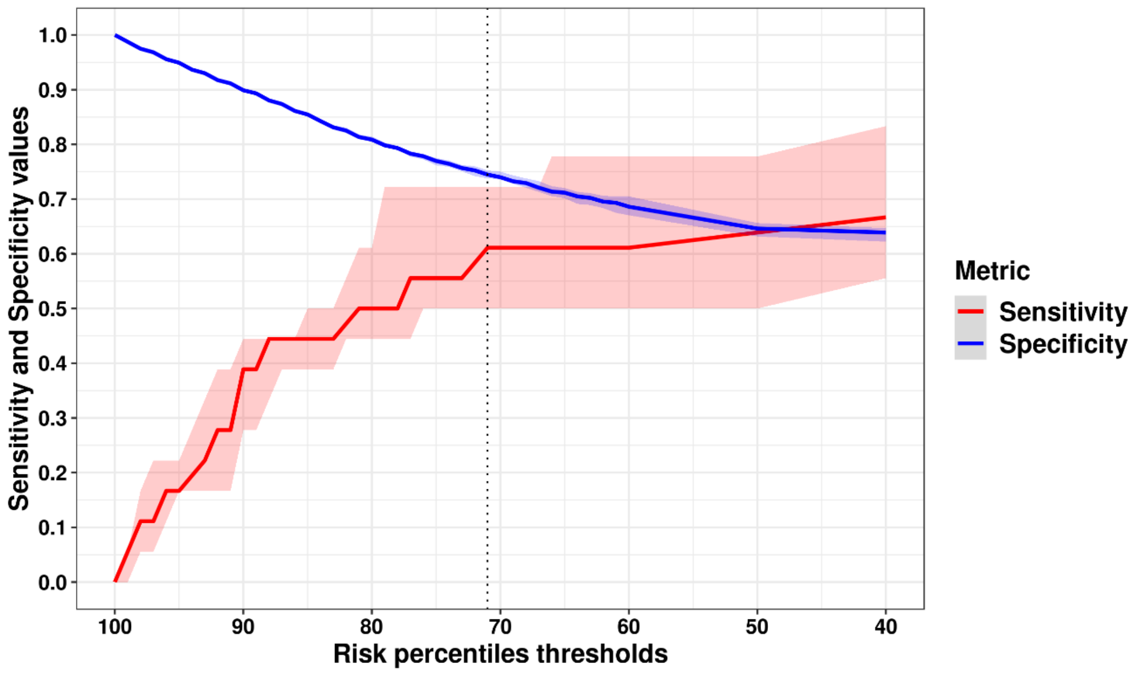


**Figure S3.** The model sensitivity and specificity. In the y-axis the estimated sensitive and specificity values and in the x-axis the risk percentiles (r). The dotted black line represents a parsimony threshold (visually chosen by the author), in which the estimated median sensitivity was 62%.

Based in our modelling results, at an r = 72$^{th}$ the median sensitivity was 62% (Figure S3), which is a suitable threshold based on the model performance, but it utilize a considerable high number of cells to make such predictions (55 cells in average by week predicted). In the example below (Figure S4), we show a histogram that represent the risk values for each cell in the Figure 5 showed it the main narrative, here we used a more rigorous threshold, r = 90$^{th}$, to identify at-risk areas with a lower number of cells (16 cells). The model correctly predicted two of the three infected cells (66% accuracy) in that week.

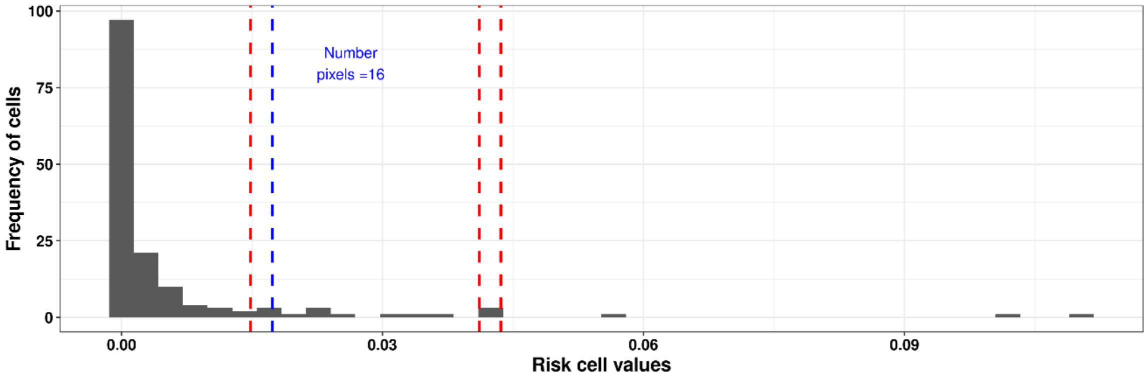


**Figure S4.** Histogram of the calculated risk values for each cell shown in Figure 5 in the main text. Horizontal blue line show the 90$^{th}$ percentile used as threshold and red lines the position of the true infected cells. Risk values generated from 100 simulations.

**Section 4: Modelling vaccination and results**

Our model assumed that vaccination modulated (reduced) the force of infection between infected and vaccinated farms to susceptible farms. The equation 1 in the main narrative becomes the following:

$\lambda net_{it} = \beta net * (N_{it} - (M_{it} * ve_{iDt}) ); \forall_i = 1, \dots, n\ suceptible\ farms; \forall_D = 1, \dots, n\ infected\ and\ vaccinated\ farms,$

where $N_i$ is the number of incoming contacts received from infected farms (undetected and detected) at the time step *t (one week)* as described earlier, but the contribution of the subset of infected farms that are vaccinated, $M$ is decreased proportionally to vaccine efficacy *ve*. Similarly, local transmission is modified (equation 4 in the mains text), as the attraction force in infected and vaccinated farms decreases proportional to vaccine efficacy, such that:

$$\lambda local_{it} = \beta local * \sum_{j=1}^{n} \left(g_{ijt} - \left(g_{ijt} * ve_{iDt}\right)\right);$$

$\forall_i = 1, \dots, n\ suceptible\ farms\ ; \forall_j = 1, \dots, n\ infected\ farms; \forall_D = 1, \dots, n\ infected\ and\ vaccinated\ farms,$

On the other hand, susceptible vaccinated farms are less likely to receive the infection, so that

$$\lambda_{it} = \lambda_{it} - (\lambda_{it} * ve_{it})$$

**The proportion of infected farms reduced for each vaccination strategy and complementary interventions**

**Table S2.** The effect of complementary interventions on the proportion of case reduction, for each farm type and for the full range of vaccine efficacies for the prophylactic #1 control strategy.

| | | **The % of infected farms reduction (sow)** | **The % of infected farms reduction (nursery)** | **The % of infected farms reduction (GDU)** | **The % of infected farms reduction (finisher)** |
|---|---|---|---|---|---|
| **Vaccine efficacy** | 0% | **Reference** | **Reference** | **Reference** | **Reference** |
| | 1% | 7 (3.5,10.5)* | 2.5 (0.3,4.7)* | 0.9 (-3.3,5.2) | 4 (1.2,6.7)* |
| | 2% | 6.1 (2.6,9.7)* | 5.4 (3.2,7.6)* | 2.5 (-1.8,6.8) | 6.9 (4.2,9.6)* |
| | 3% | 5.5 (1.9,9)* | 8.1 (5.9,10.3)* | 3 (-1.3,7.3) | 10.2 (7.5,12.9)* |
| | 4% | 5.5 (2,9)* | 10.3 (8.1,12.5)* | 1.8 (-2.5,6.1) | 13.3 (10.6,16.1)* |
| | 5% | 6 (2.5,9.6)* | 11 (8.8,13.2)* | 3.2 (-1,7.5) | 15.5 (12.7,18.2)* |
| | 10% | 10.9 (7.4,14.4)* | 18.8 (16.6,21)* | 5.1 (0.8,9.4)* | 25.3 (22.5,28)* |
| | 15% | 10.2 (6.7,13.8)* | 23.3 (21.1,25.5)* | 3.6 (-0.7,7.9) | 27.5 (24.8,30.2)* |
| | 20% | 10.6 (7.1,14.1)* | 27.2 (25,29.4)* | 3.6 (-0.7,7.9) | 30.7 (27.9,33.4)* |
| | 30% | 11.4 (7.9,14.9)* | 34.8 (32.6,37)* | 5.8 (1.5,10)* | 35 (32.3,37.7)* |
| | 50% | 12.4 (8.9,16)* | 49.9 (47.7,52.1)* | 7.1 (2.9,11.4)* | 41.4 (38.7,44.2)* |
| | 80% | 9.9 (6.4,13.5)* | 72.5 (70.3,74.7)* | 5.3 (1,9.6)* | 48.6 (45.9,51.4)* |
| | 90% | 12.1 (8.5,15.6)* | 81.6 (79.4,83.8)* | 6.9 (2.7,11.2)* | 49.5 (46.8,52.2)* |
| **Delayed time to vaccinate after pig placement** | 3 week | **Reference** | **Reference** | **Reference** | **Reference** |
| | 2 week | -0.2 (-2.1,1.8) | 0.1 (-1.2,1.3) | -1.1 (-3.5,1.3) | 0.1 (-1.4,1.6) |
| | 1 week | -0.3 (-2.3,1.6) | 1.5 (0.3,2.7)* | 0.3 (-2.1,2.7) | 2.1 (0.6,3.6)* |
| | same week | 0.1 (-1.8,2.1) | 2.1 (0.9,3.3)* | 0.5 (-1.8,2.9) | 1.4 (-0.2,2.9)* |

*$p < 0.05$.

**Table S3.** The effect of complementary interventions on the proportion of case reduction, for each farm type and for the full range of vaccine efficacies for prophylactic #2 control strategy.

| | | **The % of infected farms reduction (sow)** | **The % of infected farms reduction (nursery)** | **The % of infected farms reduction (GDU)** | **The % of infected farms reduction (finisher)** |
|---|---|---|---|---|---|
| **Vaccine efficacy** | 0% | **Reference** | **Reference** | **Reference** | **Reference** |
| | 1% | 2 (-1.5,5.4) | 5.6 (3.4,7.8)* | -7.9 (-12.7,0) | 10.8 (8.6,13)* |
| | 2% | 7.4 (3.9,10.8)* | 10 (7.8,12.2)* | 3.3 (-1.5,8.1) | 19.1 (17,21.3)* |
| | 3% | 8.4 (5,11.9)* | 15.7 (13.5,18)* | 4 (-0.8,8.7) | 26.4 (24.2,28.6)* |
| | 4% | 10.8 (7.3,14.2)* | 17.3 (15.1,19.5)* | 2.8 (-2,7.6) | 34.3 (32.1,36.5)* |
| | 5% | 13.3 (9.9,16.8)* | 19.9 (17.7,22.1)* | 4.2 (-0.6,9) | 37.1 (34.9,39.3)* |
| | 10% | 17.6 (14.2,21)* | 28 (25.7,30.2)* | 8.1 (3.3,12.9)* | 53.6 (51.4,55.8)* |
| | 15% | 18.4 (14.9,22)* | 32.5 (30.2,34.7)* | 9.1 (4.3,13.9)* | 60 (57.8,62.2)* |
| | 20% | 19.7 (16.3,23)* | 35.1 (32.9,37.3)* | 12 (7.3,16.8)* | 65.6 (63.4,67.8)* |
| | 30% | 20 (16.6,23.5)* | 42.5 (40.2,44.7)* | 12.5 (7.7,17.3)* | 71.3 (69.1,73.5)* |
| | 50% | 20.6 (17.1,24)* | 56.8 (54.5,59)* | 16.3 (11.5,21.1)* | 79.5 (77.3,81.7)* |
| | 80% | 20.8 (17.4,24)* | 76.8 (74.5,79)* | 17.2 (12.4,21.9)* | 87.4 (85.2,89.6)* |
| | 90% | 20.7 (17.3,24)* | 84.6 (82.4,86.9)* | 16.2 (11.4,21)* | 90 (87.8,92.2)* |
| **Delayed time to vaccinate after pig placement** | 3 week | **Reference** | **Reference** | **Reference** | **Reference** |
| | 2 week | 1.3 (-0.7,3.2) | 0.5 (-0.7,1.8) | 0.5 (-2.1,3.2) | 2.2 (1,3.4)* |
| | 1 week | -0.2 (-2.1,1.7) | 1.2 (0,2.5) | -0.4 (-3.1,2.3) | 3.1 (1.8,4.3)* |
| | same week | 1.2 (-0.7,3.1) | 2.1 (0.8,3.3)* | 2.1 (-0.6,4.7) | 4.4 (3.2,5.7)* |

*$p < 0.05$.

**Table S4.** The effect of complementary interventions on the proportion of case reduction, for each farm type and for the full range of vaccine efficacies for the reactive #1 control strategy.

| | | **The % of infected farms reduction (sow)** | **The % of infected farms reduction (nursery)** | **The % of infected farms reduction (GDU)** | **The % of infected farms reduction (finisher)** |
|---|---|---|---|---|---|
| **Vaccine efficacy** | 0% | **Reference** | **Reference** | **Reference** | **Reference** |
| | 1% | 4.2 (2.6,5.9)* | 0.7 (-0.6,2) | 6.2 (4.5,8)* | 2.1 (0.6,3.6)* |
| | 2% | 8.8 (7.2,10.5)* | 3.3 (2,4.5)* | 14.9 (13.1,16.6)* | 5.1 (3.6,6.6)* |
| | 3% | 12.1 (10.5,14)* | 4.9 (3.7,6.2)* | 18.3 (16.6,20)* | 7.7 (6.2,9.2)* |
| | 4% | 16.3 (14.7,18)* | 6.6 (5.4,7.9)* | 27.6 (25.8,29.3)* | 9.2 (7.7,10.7)* |
| | 5% | 18.6 (16.9,20)* | 7.6 (6.3,8.9)* | 31.9 (30.2,33.7)* | 11.9 (10.4,13.4)* |
| | 10% | 32.6 (30.9,34)* | 14.7 (13.5,16)* | 54.6 (52.8,56.3)* | 17.6 (16.1,19.1)* |
| | 15% | 39.3 (37.7,41)* | 19 (17.7,20.2)* | 67 (65.2,68.7)* | 21.2 (19.7,22.7)* |
| | 20% | 43.8 (42.1,45)* | 22.6 (21.3,23.8)* | 73.4 (71.7,75.2)* | 22.7 (21.3,24.2)* |
| | 30% | 50 (48.3,51.6)* | 29.8 (28.5,31.1)* | 79.6 (77.8,81.3)* | 26.1 (24.7,27.6)* |
| | 50% | 56.9 (55.3,58)* | 44.4 (43.1,45.6)* | 84.9 (83.1,86.6)* | 36 (34.5,37.5)* |
| | 80% | 67.3 (65.6,69)* | 68.3 (67,69.5)* | 86.9 (85.2,88.7)* | 47.6 (46.1,49.1)* |
| | 90% | 71.3 (69.6,73)* | 77.2 (75.9,78.4)* | 87.3 (85.6,89)* | 52.8 (51.3,54.3)* |
| **Average delay time for PRRSV detection** | 3 week | **Reference** | **Reference** | **Reference** | **Reference** |
| | 2 week | 0.6 (-0.3,1.5) | 0.6 (-0.1,1.3) | 1.4 (0.4,2.4)* | 0.5 (-0.4,1.3) |
| | 1 week | 1.4 (0.5,2.3)* | 0.8 (0.1,1.5)* | 2.2 (1.2,3.2)* | 0.7 (-0.1,1.5) |
| | Same week | 1.4 (0.5,2.3)* | 0.8 (0.1,1.5)* | 2.9 (1.9,3.9)* | 0.6 (-0.2,1.4) |
| **Delayed time for vaccination after PRRSV detection** | 3 week | **Reference** | **Reference** | **Reference** | **Reference** |
| | 2 week | 0.6 (-0.3,1.5) | 0.6 (-0.1,1.3) | -0.1 (-1.1,0.8) | 0.6 (-0.3,1.4) |
| | 1 week | 1 (0.1,1.9)* | 0.6 (-0.1,1.3) | 0.6 (-0.4,1.5) | -0.1 (-0.9,0.8) |
| | Same week | 0.7 (-0.2,1.6)* | 1 (0.3,1.7)* | 1.2 (0.2,2.2)* | 0.4 (-0.4,1.2) |

[1] Probability of virus detection at nursery, finisher and gilt development units (GDU) farms. For sow farms the probability was assumed to be at 95%. *p < 0.05.

**Table S5.** The effect of complementary interventions on the proportion of case reduction, for each farm type and for the full range of vaccine efficacies for the reactive #2 control strategy.

| | | **The % of infected farms reduction (sow)** | **The % of infected farms reduction (nursery)** | **The % of infected farms reduction (GDU)** | **The % of infected farms reduction (finisher)** |
|---|---|---|---|---|---|
| **Vaccine efficacy** | 0% | **Reference** | **Reference** | **Reference** | **Reference** |
| | 1% | 6 (5.1,6.9)* | 4.5 (3.8,5.2)* | 7.7 (6.7,8.7)* | 8.4 (7.6,9.1)* |
| | 2% | 11.8 (10.9,13)* | 8.3 (7.7,9)* | 15.4 (14.4,16.3)* | 15.3 (14.6,16.1)* |
| | 3% | 16.3 (15.4,17)* | 10.4 (9.7,11.1)* | 21.8 (20.8,22.8)* | 21.1 (20.3,21.8)* |
| | 4% | 20.8 (19.9,22)* | 13.5 (12.8,14.2)* | 29.3 (28.3,30.3)* | 26.8 (26.1,27.6)* |
| | 5% | 25.2 (24.3,26)* | 16.7 (16,17.4)* | 36.5 (35.6,37.5)* | 32.1 (31.4,32.9)* |
| | 10% | 41.8 (40.8,43)* | 26.1 (25.4,26.8)* | 61 (60,62)* | 49.7 (48.9,50.4)* |
| | 15% | 50.5 (49.6,52)* | 31.8 (31.1,32.4)* | 73.9 (72.9,74.9)* | 59.5 (58.8,60.2)* |
| | 20% | 56.4 (55.4,57)* | 37.4 (36.7,38.1)* | 80.9 (79.9,81.9)* | 65.4 (64.7,66.1)* |
| | 30% | 62.6 (61.7,63.5) | 45.2 (44.5,45.9)* | 87.3 (86.4,88.3)* | 72.2 (71.4,72.9)* |
| | 50% | 71.6 (70.7,73)* | 60.2 (59.5,60.9)* | 91.9 (91,92.9)* | 81.2 (80.5,81.9)* |
| | 80% | 84.1 (83.2,85)* | 82.9 (82.2,83.6)* | 94 (93.1,95)* | 92.8 (92,93.5)* |
| | 90% | 89.3 (88.3,90)* | 90.8 (90.1,91.5)* | 94.6 (93.6,95.6)* | 96.4 (95.7,97.1)* |
| **Average delay time for PRRSV detection** | 3 week | **Reference** | **Reference** | **Reference** | **Reference** |
| | 2 week | 0.2 (-0.4,0.7) | 0.2 (-0.2,0.6) | 0.3 (-0.2,0.9) | 0.7 (0.3,1.1)* |
| | 1 week | 0.3 (-0.2,0.8) | 0.3 (-0.1,0.7) | 0.6 (0.1,1.2)* | 0.8 (0.4,1.3)* |
| | Same week | 0.9 (0.4,1.4)* | 0.6 (0.3,1)* | 0.4 (-0.1,1) | 1.2 (0.7,1.6)* |
| **Delayed time for vaccination after PRRSV detection** | 3 week | **Reference** | **Reference** | **Reference** | **Reference** |
| | 2 week | 0.8 (0.3,1.4)* | 0.8 (0.4,1.2)* | 1.3 (0.8,1.9)* | 1.5 (1.1,1.9)* |
| | 1 week | 1.8 (1.3,2.3)* | 1.3 (0.9,1.7)* | 2.2 (1.7,2.7)* | 2.5 (2.1,2.9)* |
| | Same week | 1.8 (1.3,2.3)* | 1.6 (1.2,2)* | 2.4 (1.8,2.9)* | 3.3 (2.8,3.7)* |
| **Prob. of PRRSV detection[1]** | 7% | **Reference** | **Reference** | **Reference** | **Reference** |
| | 50% | 0.9 (0.4,1.3)* | 1.7 (1.4,2.1)* | 0.2 (-0.2,0.7) | 3.6 (3.3,4)* |
| | 95% | 0.9 (0.4,1.3)* | 2.3 (2,2.6)* | 0.5 (0.1,1)* | 4.9 (4.5,5.2)* |

[1] Probability of virus detection at nursery, finisher and gilt development units (GDU) farms. For sow farms the probability was assumed to be at 95%. *$p < 0.05$.

**Table S6.** The effect of complementary interventions on the proportion of case reduction, for each farm type and for the full range of vaccine efficacies for combined control strategy.

| | | **The % of infected farms reduction (sow)** | **The % of infected farms reduction (nursery)** | **The % of infected farms reduction (GDU)** | **The % of infected farms reduction (finisher)** |
|---|---|---|---|---|---|
| **Vaccine efficacy** | 0% | **Reference** | **Reference** | **Reference** | **Reference** |
| | 1% | 8.3 (7.9,8.8)* | 8.2 (7.9,8.5)* | 9.1 (8.7,9.6)* | 15.5 (15.2,15.7)* |
| | 2% | 15.1 (14.7,16)* | 14.7 (14.4,14.9)* | 17.1 (16.7,17.6)* | 28.1 (27.8,28.4)* |
| | 3% | 21.4 (21,21.8)* | 20.1 (19.8,20.4)* | 25.1 (24.7,25.5)* | 38.4 (38.2,38.7)* |
| | 4% | 26.6 (26.2,27)* | 24.5 (24.2,24.8)* | 32.3 (31.9,32.8)* | 46.7 (46.4,46.9)* |
| | 5% | 31.4 (31,31.8)* | 28.3 (28,28.5)* | 38.9 (38.5,39.4)* | 53.7 (53.4,53.9)* |
| | 10% | 47.6 (47.2,48)* | 41 (40.8,41.3)* | 63 (62.5,63.4)* | 72.9 (72.7,73.2)* |
| | 15% | 56 (55.6,56.4)* | 48.7 (48.4,49)* | 76.4 (75.9,76.8)* | 81 (80.7,81.3)* |
| | 20% | 60.9 (60.5,61)* | 54.5 (54.2,54.8)* | 83.2 (82.8,83.6)* | 85.2 (84.9,85.4)* |
| | 30% | 66.4 (65.9,67)* | 64.7 (64.4,65)* | 89 (88.5,89.4)* | 89.9 (89.7,90.2)* |
| | 50% | 73.8 (73.4,74)* | 80.9 (80.6,81.2)* | 93.1 (92.7,93.6)* | 95.2 (94.9,95.4)* |
| | 80% | 85 (84.6,85.4)* | 95.5 (95.2,95.7)* | 94.4 (94,94.9)* | 98.8 (98.5,99.1)* |
| | 90% | 89.6 (89.2,90)* | 98 (97.7,98.3)* | 94.6 (94.2,95.1)* | 99.5 (99.2,99.8)* |
| **Delayed time to vaccinate after pig placement** | 3 week | **Reference** | **Reference** | **Reference** | **Reference** |
| | 2 week | 0.2 (0,0.4) | 0.3 (0.2,0.5)* | 0.2 (-0.1,0.4) | 0.5 (0.4,0.7)* |
| | 1 week | 0 (-0.2,0.2) | 0.7 (0.5,0.9)* | 0.2 (-0.1,0.4) | 1.2 (1,1.3)* |
| | Same week | 0.3 (0.1,0.6)* | 1.3 (1.1,1.4)* | 0.3 (0,0.5)* | 2 (1.9,2.2)* |
| **Average delay time for PRRSV detection** | 3 week | **Reference** | **Reference** | **Reference** | **Reference** |
| | 2 week | 0 (-0.3,0.2) | 0 (-0.1,0.2) | 0 (-0.3,0.2) | 0.1 (-0.1,0.2) |
| | 1 week | 0 (-0.2,0.2) | 0.1 (0,0.3) | 0.3 (0,0.5)* | 0 (-0.1,0.2) |
| | Same week | 0 (-0.2,0.3) | 0.1 (-0.1,0.2) | 0.5 (0.2,0.7)* | 0 (-0.1,0.2) |
| **Delayed time for vaccinatio n after** | 3 week | **Reference** | **Reference** | **Reference** | **Reference** |
| | 2 week | 0.7 (0.5,1)* | 0.2 (0.1,0.4)* | 1.4 (1.2,1.7)* | 0.4 (0.3,0.6)* |
| | 1 week | 1.4 (1.2,1.6)* | 0.5 (0.4,0.7)* | 2.3 (2,2.5)* | 0.8 (0.7,1)* |

| | | | | | |
|---|---|---|---|---|---|
| **PRRSV detection** | Same week | 1.4 (1.2,1.6)* | 0.4 (0.3,0.6)* | 2.3 (2.1,2.6)* | 0.9 (0.8,1.1)* |
| **Prob. of PRRSV detection[1]** | 7% | **Reference** | **Reference** | **Reference** | **Reference** |
| | 50% | 0.1 (-0.1,0.3) | 0.2 (0,0.3)* | -0.1 (-0.4,0.1) | 0.2 (0.1,0.3)* |
| | 95% | 0.4 (0.2,0.5)* | 0.3 (0.2,0.4)* | 0.1 (-0.1,0.3) | 0.3 (0.2,0.4)* |

[1] Probability of virus detection at nursery, finisher and gilt development units (GDU) farms. For sow farms the probability was assumed to be at 95%. *$p < 0.05$.

**Section 5: Summarized network analysis of between-farm movements**

**Table S7.** Number of in-going and outgoing contacts by production types from June 01, 2019 to December 05, 2019. Multiple movements between two farms in the same date were grouped and considered as one contact.

| Farm type | In-going movements | Out-going movements |
|---|---|---|
| **Sow** | 4.375 | 13.822 |
| **Nursery** | 11.860 | 7.905 |
| **Gilt development unit** | 881 | 3.488 |
| **Finisher** | 8.558 | 456* |

***** Movements to slaughter houses or markets were removed.

**Section 6: The density of farms within 5 km, 10 km and 20 km as a proxy for neighborhood contacts**

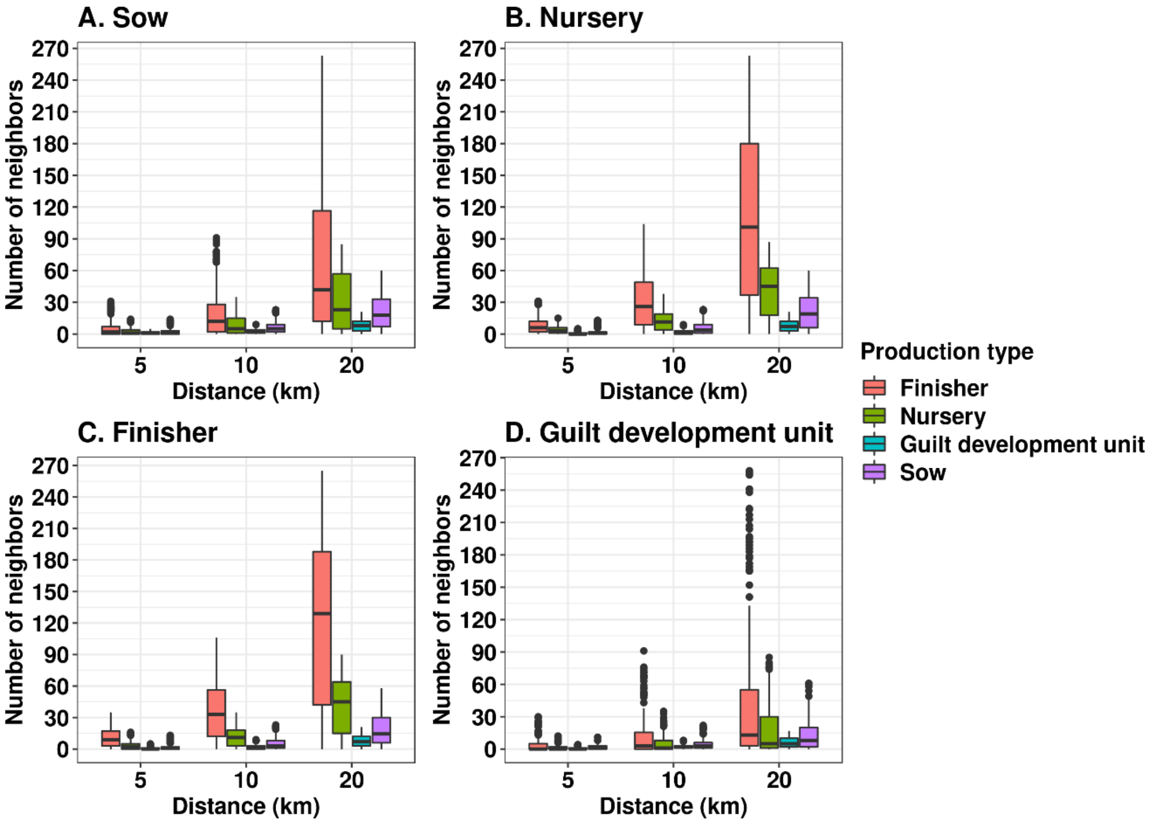


**Figure S5.** Number of farm neighbors in different radius distances from **A**) Sow farms, **B**) Nursery farms, **C**) Finisher farms and **D**) Gilt development units.

Using Kruskal-Wallis and Dunn's post-hoc test we evaluated the differences of the number of neighbors within each radius size. For example, for a radius of 5 km, finisher farms were the most frequent neighbor of all other farm types ($p < 0.05$). Also, this pattern seems to be linear as the size of neighborhoods increased (Figure S7). We tested if the median number of finisher neighbors was higher in finisher farms compared with nursery farms, the result showed that nursery farms had a median of 6 finisher farm neighbors, a significant lower median than finisher farms which was 9 (Wilcoxon test $p < 0.05$).
